# Polysialic acid is a versatile marker for retinal Müller glia in common vertebrate model organisms and systems

**DOI:** 10.1101/2025.09.04.672712

**Authors:** A. Nicol San Juan, Allison D. N. N. Tran, Debarati Hore, Linnea M. Kriese, Natalie H. N. Van Engelen, Maria Sharkova, Nicole C. L. Noel, Carmanah Hunter, Olga Spencer Carreira, Jennifer C. Hocking, Lisa M. Willis, Brittany J. Carr

**Author notes:** Corresponding Author: Brittany J. Carr. **Conflict of Interest Disclosure:** The authors declare no conflict of interest.

## Abstract

Müller glia are retinal support cells that play crucial roles in tissue structure, waste management, and repair. A challenge for the field has been to find Müller glia markers that detect non-reactive cells or work well in non-mammalian models. We introduce two novel markers for identifying reactive and non-reactive Müller glia in vertebrate retinas: a modified enzyme lectin (GFP-EndoN_DM_) and a monoclonal antibody (mAb735). These markers recognize polysialic acid (polySia), which is a highly conserved glycosylation modification in humans and vertebrates. In the retina, polySia is present on Müller glia predominantly in the form of polySia-NCAM. We used GFP-EndoN_DM_ and mAb735 to investigate polySia distribution in Müller glia of fish, amphibians, reptiles, birds, rodents, retinal organoids, and humans. In adult retinas of most species, polySia was localized to the Müller glia and spanned outer to inner retina, with inner plexiform layer (IPL) sublaminae ramifications. Gliosis was also detectable in degenerating murine retinas. Notable species differences were that only outer retinal regions of Müller glia were labelled in adult zebrafish, whereas the outer Müller glia body up to the first IPL sublamina was labelled in adult turquoise killifish. There was no significant retinal polySia labeling in larval zebrafish, but it was present in the brain. Larval turquoise killifish have polySia throughout the retina, similar to other adult vertebrates. Labelling polySia expands the scientific toolbox for Müller glia markers, and offers a versatile way to visualize and monitor structural changes in non-reactive and reactive Müller glia across most vertebrate species.

**Highlights:**

- PolySia is a highly conserved Müller glia marker in vertebrate retina and retinal organoids
- Labeling Müller glia with markers of polySia, facilitates the analysis of fine Müller glia processes in the IPL sublaminae, whole Müller glia morphology, and degenerative gliosis
- PolySia can be labeled with an engineered lectin conjugate (GFP-EndoN_DM_) and a commercially-available monoclonal antibody

## **1.** Introduction

### 1.1 Müller glia

Müller glia are the principal macroglial cells in the vertebrate retina. They are homologous to the radial glia of the brain and spinal cord, and are integral supporters of retinal architecture, metabolism, and homeostasis. Müller glia have many functions including water and ion regulation^1^, retinoid metabolism and visual pigment recycling^2^, and energy production^3^. In addition, Müller glia also participate in phagocytosis and retinal waste removal, both during development^4^ and in response to retinal degenerative disease^5^. Although they are one of the last cell types to differentiate during retinal development^6^, Müller glia are essential in guiding retinal organization and maintaining retinal laminations by supporting retinal plasticity, neuronal pruning, and retinal information processing^1,4,7,8^. Selective destruction of the Müller glia results in retinal dysplasia, photoreceptor death, retinal vascular abnormalities, and retinal pigment epithelium (RPE) dysfunction^9–11^. From apical to basal retina, Müller glia occupy nearly the entire neural retinal space, and interact with or comprise significant structural components of the retina, including the outer segment layer (OS), the external limiting membrane (ELM), the outer nuclear layer (ONL), the outer plexiform layer (OPL), the inner nuclear layer (INL), the inner plexiform layer (IPL), the ganglion cell layer (GCL), the nerve fibre layer (NFL), and the internal limiting membrane (ILM). The Müller glia apical processes, which extend beyond the ELM into the OS space, may interact with RPE microvilli to aid in cone stability or chromophore metabolism and recycling^12^. Müller glia interact with outer and inner retinal neurons in both plexiform layers, and contribute to retinal homeostasis by supporting the neurons with trophic molecules, removing waste, ion regulation, neurotransmitter release, and regulation of ATP^13^. Müller glia contribute to retinal integrity not only by forming connections with retinal neurons^14^, but also by formation of the ELM and the ILM^15^. The ELM and ILM are not true membranes, but instead comprise tight and adherent junctions between adjacent Müller glia and the photoreceptors, which form a type of semi-permeable membrane that can regulate molecule transport and add structural support to the retina^16,17^. The ELM is formed as a junction with the photoreceptor inner segments, whereas the ILM is formed by the Müller glia endfeet at the vitreoretinal interface. Finally, in some species like fish and amphibians, Müller glia support retinal regeneration and repair by de-differentiating into retinal progenitors, acting as stem cells to re-populate retinal neurons lost to injury^18–20^.

In addition to important roles in retinal homeostasis, Müller glia become reactive in response to retinal degenerative pathology by undergoing morphological, physiological, and biochemical changes^21,22^. These gliotic alterations are intended to protect the retina, by releasing neurotropic factors, scavenging free radicals, modulating glutamate uptake, and facilitating the formation of glial scars, which isolate diseased tissue from healthy tissues^21–23^. Sometimes, Müller glia can instead be harmful, particularly by contributing to the development of edema and preventing retinal repair and remodeling as a result of glial scarring^24–27^. Two well known indicators of Müller glial reactivity and gliosis are the upregulation of the intermediate filament protein, glial fibrillary acidic protein (GFAP) and changes in glutamine synthetase (GS), a Müller cell-specific enzyme normally involved in neurotransmitter recycling^28–30^. Consequently, these two Müller glia markers are the most popular for use in biomedical research. GFAP and GS are less-than-ideal tools for revealing the morphology of the entire Müller glia cell, however, especially the fine processes in the plexiform layers^31,32^. They are also not very effective markers for Müller glia under homeostatic conditions, because they sometimes do not have significant expression of GFAP or GS^33^. Thus, there is significant room for improvement in the development or characterization of a versatile Müller glia marker for use across various stages of retinal health, and in different species.

### 1.2 PolySia-NCAM

Glycosylation is the post-translational process of covalently attaching carbohydrates to a target protein or macromolecule, with virtually all cell surface proteins undergoing modification with glycans to form a glycocalyx around the cell membrane^34^. Protein glycosylation increases proteomic diversity, provides biochemical stability, and, in some cases, can confer additional functions to proteins. The glycocalyx on the outside of the cell, which can be up to 500 nm thick, is involved with cell-cell and cell-extracellular matrix communication, cell migration and adhesion, and pathogen-host interactions^35,36^. Sialic acids or neuraminic acids comprise a family of 9-carbon carboxylated sugars, named 3-deoxy-D-*glycero*-D-*galacto*-non-2-ulosonic acids. *N*-acetylneuraminic acid (Neu5Ac) and *N*-glycolylneuraminic acid (Neu5Gc) are the most common sialic acids in vertebrates and differ from one another only in the presence of an additional hydroxyl group in Neu5Gc. In mammals, the Neu5Gc modification is created via hydroxylation of Neu5Ac by the cytidine monophosphate-N-acetylneuraminic acid hydroxylase (CMAH) enzyme^37^. Certain species, including humans, deer, chickens, and brown bats, have lost the ability to make Neu5Gc due to dysfunctional or non-functional CMAH^38^. Regardless of whether a species can make Neu5Gc, the dominant sialic acid in the brain and retina is almost exclusively Neu5Ac^39^. These sialic acids frequently occupy the terminal position on various glycoproteins and glycolipids in mono-, oligo-, or polymeric forms^34^. Polysialic acid (polySia) is a is negatively charged linear homopolymer of 8 or more sialic acids which, in the vertebrate nervous system, are connected though α2,8-linkages. PolySia is reported to modify properties of cell-cell associations, including signaling, adhesion, and migration^40^. It also plays a significant role in cancer metastasis, and has been investigated in the context of cancer vaccines^41,42^.

In the central nervous system, polySia is found bound primarily to the membrane protein neural cell adhesion molecule (NCAM)^43,44^. NCAM is found in neurons, glial cells, and Schwann cells^45–47^, and can be post-translationally modified with polySia to form polySia-NCAM^48,49^. It is thought that polySia-NCAM reduces the strength of the adhesion forces between cells by steric hindrance, making it harder for cells come into contact and decreasing cellular adhesion^48,50^. This decreased adhesion property of polySia-NCAM has led to its implication in the regulation of cell migration, synapse formation, and neuronal plasticity^51^. Correspondingly, polySia expression in the brain is high during embryonic development^44,52,53^, but it also remains present at lower expression levels in adults, particularly in areas important for neuro-plasticity or neuro-progenitor populations^54–58^. Retinal expression of polySia-NCAM is less well studied. NCAM alone has been reported to be expressed in adult rodent Müller glia and putative optic nerve astrocytes, and the inner and outer plexiform layers in bovine and frog retina^52,59–61^. Retinal polySia expression has been investigated previously in the rat, salamander and zebrafish^60,62,63^. Interestingly, only the rat paper identified the polySia to be localized to the Müller glia. In salamander and zebrafish, polySia expression was reported to be localized primarily to the ONL, near the cone photoreceptor cell bodies. In the salamander, polySia expression was also reported within the INL, IPL, GCL, and NFL^62^.

The high conservation of polySia structure and its localization to the cell membrane makes it an attractive target for labeling Müller glia, enabling the study of glial morphology and behavior in the vertebrate retina. We found that polySia is an effective and versatile marker for Müller glia in a wide variety of vertebrate species, from amphibians to humans, as well as in retinal organoids. We also found notable differences in polySia localization on the Müller glia of fishes compared to all other vertebrates. Contrary to reports of significant differences in polySia expression during development in the brain, there is a less apparent change in polySia expression in retinal Müller glia expression of polySia between embryonic and adult retinas. The exception is again fishes; polySia is absent in larval zebrafish larval and less confined in larval killifish compared to adult killifish.

## 2. Materials and Methods

### 2.1 Human and Animal Ethics Statements

Human post-mortem eye tissues were obtained through collaborators under approved protocols. Retinal tissues were donated from 3 men, aged late-sixties to early eighties, with no known prior history of ocular disease (Lions Eye Bank, University of Iowa). The use of these tissues for this study was approved by the University of Alberta (Pro00147177), and carried out in accordance with the Health Canada and Public Health Agency of Canada (PHAC) Research Ethics Board (REB), guided by the principles of the second edition of the Tri-Council Policy Statement: Ethical Conduct for Research Involving Humans (TCPS 2). This study was performed in accordance with the ethical standards as laid down in the 1964 Declaration of Helsinki and its later amendments.

Animal experiments were approved by the Canadian Council on Animal Care (CCAC) and Animal Care and Use Committees (ACUC). We examined adult retinas of 8 different vertebrate non-human species: African turquoise killifish (*Nothobranchius furzeri*), zebrafish (*Danio rerio*), African clawed frog (*Xenopus laevis*), hummingbird (*Calypte anna*), fire skink (*Lepidothyris fernandi*), zebra finch (*Taeniopygia guttata*), mouse (*Mus musculus*), and rat (*Rattus norvegicus*). Adult frog eyes were obtained under the University of Alberta approved animal ethics protocol (AUP00004203) and adult zebrafish eyes were obtained under the University of Alberta approved ethics protocol (AUP00001476). Killifish experimental procedures were conducted in accordance with the UK Home Office Animals (Scientific Procedures) Act of 1986 and killifish eyes were obtained under the animal licenses 70/7391 and PP2133797.

All other animal tissues were donated to the Carr lab from various researchers after the animals were euthanized by appropriate personnel under appropriate protocols (mouse, rat) or obtained under appropriate wildlife permits and approved protocols (fire skink, hummingbird, zebra finch). We also investigated two cephalopod species. Octopus (*Octopus bimaculoides*) and squid (*Euprymna berryi*) were acquired from the Cephalopod Resource Center (Woods Hole, MA, USA) and kept at the University of Oregon, until fixed tissues were sent to the Carr lab. Cephalopod studies were conducted with approved protocols from the University of Oregon Animal Care Services, in compliance with the Association for Assessment and Accreditation of Laboratory Animal Care International guidelines. Animal husbandry and protocols were carried out in accordance with published guidelines for the care and welfare of cephalopods in the laboratory^64^. Figures are representative images from an n = 3-4 for all species, except for the fire skink (n = 1), zebra finch (n = 2), octopus (n = 1), and squid (n = 1).

Most tissues were from wildtype animals. Exceptions were 2/3 of the adult Sprague Dawley rats, which were Thy1-GFP transgenics, the postnatal rats were culls from a connexion (Cx) 40 heterozygous breeding (25% Cx^+/+^, 25% Cx40^-/-^, 50% Cx40^+/-^), and the postnatal mice, which were floxed R26^lsl^-Cmah mice^65^. Thy1 is a surface glycosylphosphatidylinositol-anchored protein with proposed roles in the control of cellular differentiation, cell adhesion, cell signalling, and immunity^66,67^. In the adult retina, THY1 protein is reported to be expressed in a subset of retinal amacrine and ganglion cells^68–70^, which we confirmed in this study. The presence of Thy1-GFP did not obviously affect retinal polySia expression patterns compared to the wildtype rat (**Supplementary Fig. S1**). Cx40 is associated with the development of retinal vasculature^71^, and we did not observe any significant differences in retinal vasculature or retinal labeling patterns between the postnatal rats investigated in this study (n = 6). Dams of postnatal mice were not fed tamoxifen during pregnancy, so the floxed R26^lsl^-Cmah mice should represent a wildtype phenotype. We observed no differences in the labelling patterns between littermates (n = 7).

### 2.2 Organoid Sources and Differentiation

Retinal organoids were obtained from collaborators under approved protocols (NOVA Medical School, Universidade Nova de Lisboa). Two human inducible pluripotent stem cell (hiPSC) lines were used in this study: a Parkinson’s patient commercialized cell line (ND50040, NHCDR/RUCDR) and a healthy control cell line (IMR90-4, WiCell). Cell lines were maintained and expanded to 80% confluency, after which a retinal organoid differentiation protocol was implemented as described previously^72^. Briefly, hiPSCs were lifted (Versene, Thermo Fisher Scientific) and split at 3000 cells/well in a Nunclon Sphera 96-well U-bottom plate (Thermo Fisher Scientific) using mTeSR™ Plus medium supplemented with 10 µM of ROCK inhibitor (Focus Biomolecules). The cells were grown as aggregates for 6 days as an adaptation period to Neural Induction Medium (NIM) containing DMEM/F12, 1% N2 supplement, 1L×Lnon-essential amino acids, 1L×LGlutaMax (Gibco, Thermo Fisher Scientific), and 2 mg/ml Heparin (Sigma-Aldrich). On day 6, 1.5 nM of bone morphogenic protein-4 (BMP4, Peprotech) was added to fresh NIM, and on day 7, EBs were transferred to growth factor–reduced matrigel-coated 6-well plates (Corning). The medium was replaced by half-fresh NIM on days 9, 12, and 15. On day 16, the medium was replaced by retinal differentiation medium (RDM) containing DMEM:F12 (3:1), 2% B27 minus vitamin A, 1L×Lnon-essential amino acids, 1L×LGlutaMax, 1L×Lantibiotic/antimycotic (Gibco, Thermo Fisher Scientific). This medium was changed every 2–3 days. At day 30, optic vesicles were manually dissected using a surgical scalpel (SM65A, Swann-Morton Ltd) under the microscope EVOS XL core (Thermo Fisher Scientific). Dissected organoids were maintained in suspension flasks (Sarstedt) in 3D-retinal differentiation medium (3D-RDM) containing DMEM:F12 (3:1), 2% B27 minus vitamin A, 1L×Lnon-essential amino acids, 1L×LGlutaMax, 1L×Lantibiotic/antimycotic, 5% FBS, 1:1000 chemically defined lipid supplement (Gibco, Thermo Fisher Scientific), 100 µM taurine (Sigma-Aldrich), and 1 µM of all-trans retinoic acid (RA, Sigma-Aldrich) until day 120. Fixed and cryoprotected organoid tissues (day 120) were shipped to us, and then we sectioned and labelled them according to the protocol described below.

### 2.3 Retinal Tissue Preparation and Cryosectioning

Whole eyes prepared in the lab were removed and fixed in 4% paraformaldehyde (PFA) + 3% sucrose in 0.1 M sodium phosphate buffer (PB, pH 7.4) (0.774M Na_2_HPO_4_ + 0.226M NaH_2_PO_4,_ pH 7.4) overnight at 4°C. Eyes obtained under wildlife permits (fire skink, hummingbird, zebra finch) were perfused with 4% PFA and then kept in fixative for various periods, from 4 days (hummingbird, fire skink) to 2 years (zebra finch). Cephalopod eyes were obtained by deeply anesthetizing them in artificial seawater (ASW; 460mM NaCl_2_, 10mM KCl, 10mM glucose, 10mM HEPES, 55mM MgCl_2_, 11mM CaCl_2_, 2mM glutamine, pH 7.4) supplemented with 110mM MgCl_2_, followed by rapid decapitation and fixation in 4% PFA in ASW at 4°C. Frog eyes labeled with GFAP primary antibody (**Fig. 4E**) were fixed in 80% methanol + 20% dimethylsulfoxide at -20°C (Dent’s fixative protocol) overnight since 4% PFA + 3% sucrose fixation eliminated antibody binding. After fixation, all eyes were processed the same way. Post-fixed eyes were cryoprotected in increasing gradients of sucrose (10%, 20%, and then 30%) diluted in 0.1 M PB. Fully cryoprotected eyes were then embedded in Tissue Tek O.C.T. (*S*akura Finetek, Torrance, CA, USA) and sagittal sections were cut at a thickness of 12-16 μm. Sections were picked up on room temperature SuperFrost slides (Thermo Fisher Scientific, Waltham, Massachusetts, USA) and then left on a warm (42°C) hot plate for 8-10 min to increase tissue adherence to the slides. Prepared slides were stored at -20°C until required.

### 2.4 Immunofluorescence and Epi-Fluorescence Microscopy

Frozen slides were thawed at room temperature, tissues were outlined with a hydrophobic marker (ImmEdge™, MJC Biolynx, Brockville, ON, Canada), and then slides were washed 3x8 mins in 1X phosphate buffered saline (PBS, pH 7.4) (137 mM NaCl + 2.7 mM KCl + 10 mM Na_2_HPO_4_ + 1.8 mM KH_2_PO_4_) at room temperature (∼20°C). After washing, slides were blocked in blocking buffer (1% goat serum + 0.1% Triton X-100 in 1X PBS) for 30 minutes in a humid chamber at room temperature.

Slides were washed again 3x8 min in 1X PBS and then sections were labeled with GFP-EndoN_DM_ (10 µg/mL; Dr. Lisa Willis^73^, University of Alberta) or a primary antibody to polySia, mAb735 (1 µg/mL; Absolute Antibody, AB00240-2.0) in dilution buffer (0.1% goat serum + 0.1% TX-100 in 1X PBS) overnight in a humid chamber at room temperature. After the overnight incubation, slides were washed three times for 8 min each in 1X PBS, and then counter labels and the secondary antibody for mAb735 (AF488 goat anti-mouse, Jackson Immunoresearch, West Grove, PA, USA) were applied in dilution buffer and incubated for 4–6 h at 20°C. All tissues were counter labeled with Hoechst 33342 (Millipore Sigma, Burlington, Massachusetts, USA) and wheat germ agglutinin (Thermo Fisher, Waltham, Massachusetts, USA) to visualize nuclei and photoreceptor outer segments, respectively. Slides were washed a final time for 3x8 min in 1X PBS and coverslipped using Mowiol Cover Slip Mounting Medium without DABCO^74^. For double labeling experiments, retinal sections were incubated with GFP-EndoN_DM_ and the primary antibody overnight at the same time. Antibodies used in this study and their concentrations are listed in **Supplementary Table 1**.

Slides were imaged using either a 1) Leica DM6000B epi-fluorescence microscope with a 20x objective (air, N.A. 0.50), Leica K3M Camera (3072 px x 2048 px), and Leica Software; 2) a Zeiss Airyscan LSM900 with a 40x objective (water, N.A. 1.2; killifish); or a 3) Zeiss LSM 700 with a 63x objective (Oil, N.A. 1.40; zebrafish). Post-imaging processing was performed using Affinity Photo 2 (Serif, West Bridgford, UK) and FIJI^75^.

### 2.5 GFP-EndoN_DM_ and mAb735 Characterization

Endoneuraminidase F (EndoN) is a polySia hydrolase that was originally isolated from the tailspike of an *E. coli* K1-specific bacteriophage^76,77^. A double mutant (EndoN_DM_) was subsequently engineered to be a catalytically dead lectin that can detect and bind to polySia chains with high affinity (K_D_ = 10^−8^ M) without hydrolysing it^78^. GFP-tagged EndoN_DM_ (GFP-EndoN_DM_) is an simple and efficient marker that has been used for the detection of polySia in numerous cell culture systems and applications^42,76,79,80^. mAb735 is a primary mouse monoclonal antibody (Absolute antibody, AB00240) that specifically recognizes α2,8-linked polySia chains of 8 or longer^81,82^. In adult mice, mAb735 labels polySia-NCAM, attributed to Müller glia and astrocytes in the retina and optic nerve^52^. mAb735 has also previously been reported to label Müller glia in adult rat retina^60^.

To validate that retinal GFP-EndoN_DM_ and mAb735 positive signal in our vertebrate retinas were due to binding to polySia, slides with mouse retina were incubated for 30 min at 37°C with the active polySia hydrolase EndoN^77^. PBS was used as a negative control to keep slides hydrated in the absence of the hydrolase. After the 30 min incubation, slides were washed 3x8 min with 1X PBS and then labeled with GFP-EndoN_DM_ or mAb735. GFP-EndoN_DM_ and mAb735 labeling were eliminated by pre-treatment with polySia hydrolase (**Supplementary Fig. S2B,D**). We found that there was non-specific secondary antibody labeling in the intraretinal and vitreal blood vessels, the RPE, the choroid, and the sclera in mice and rats, this is expected due to cross-reactivity of mouse secondary antibodies to rodent endogenous blood plasma IgGs (**Supplementary Fig. S2E**). This cross reactivity was not seen in non-rodent species.

## 3. Results

### 3.1 GFP-EndoN_DM_ and mAb735 localization in developing non-human vertebrate retinas

To determine the extent of polySia expression in developing non-human vertebrate retinas, we investigated the pattern of GFP-EndoN_DM_ and mAb735 labeling in the early free-swimming larval retinas of zebrafish (*Danio rerio*) (∼5 days post fertilization (dpf)), turquoise killifish (*Nothobranchius furzeri*) (golden eye stage, ∼21 dpf), and African clawed frog (*Xenopus laevis*) (N.F stage 40, ∼5 dpf)^83–85^. We also looked at postnatal (P3) mouse (*Mus musculus*) and (P1) rat (*Rattus norvegicus*) retina, before retinal lamination is complete. GFP-EndoN_DM_ and mAb735 labeling patterns were comparable (**Fig. 1**). PolySia was detected in the retina and brain of larval killifish (**Fig. 1A-B, I**) and larval frog (**Fig. 1E-F,K**). It was also detected in the retinas of postnatal P3 mouse (**Fig. 1G,L**) and postnatal P1 rat (**Fig. 1H,M)**, but not in larval zebrafish retina (**Fig. 1D,J**). Although there was no GFP-EndoN_DM_ or mAb735 signal in the larval zebrafish retina, there was significant positive GFP-EndoN_DM_ signal in the larval zebrafish brain (**Fig. 1C**). In larval frog and killifish retinas, GFP-EndoN_DM_ and mAb735 positive signal was present in the entire Müller glial cell body, from the ELM to the endfeet, but was strongest near the ONL and the OPL, and the NFL. In P1 rat and P3 mouse, the retina is not yet fully laminated, but there was a strong GFP-EndoN_DM_ and mAb735 positive band near the lower one-third of the retina (IPL) and diffuse labeling through the neuroblast layer (NBL) up to the developing ELM (**Fig. 1E,F; J,K**).

**Figure 1.**
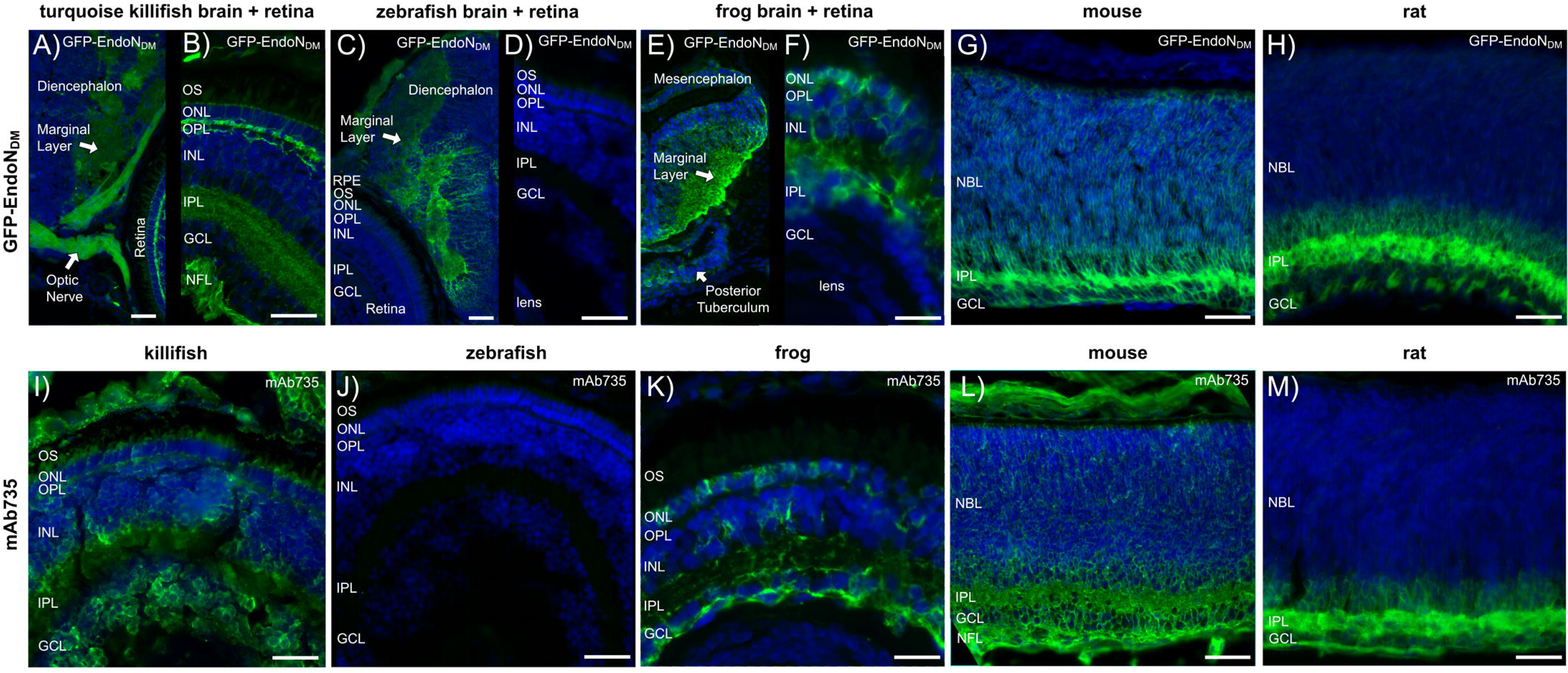
GFP-EndoN_DM_ and mAb735 labeling in developing killifish (A-B,I), zebrafish (C-D, J), frog (E-F,K), mouse (G,L), and rat (H,M). PolySia is present in the retina and brain of killifish and frog (A-B, E,F), in the brain of zebrafish (C), and in the retina of postnatal mice and rats (G,H,L,M). Minor differences in labeling, such as increased scleral reactivity observed in mAb735-labeled mice (L) are the result of cross-reactivity of the secondary antibody (Supplementary Fig. S2). *Labels*: Green, GFP-EndoN_DM_ (A-H), mAb735 (I-M); Blue, Hoechst. *Abbreviations*: OS, outer segments; ONL, outer nuclear layer; OPL, outer plexiform layer; INL, inner nuclear layer; IPL, inner plexiform layer; GCL, ganglion cell layer; NFL, nerve fiber layer; NBL, neuroblast layer. Scale bar = 50 µm. n = 3-7.

### 3.2 GFP-EndoN_DM_ and mAb735 localization in adult non-human vertebrate and cephalopod retinas

To investigate polySia expression in adult non-human vertebrate retinas, we investigated the localization of GFP-EndoN_DM_ and mAb735 in 8 different species: turquoise killifish, zebrafish, hummingbird (*Calypte anna*), zebra finch (*Taeniopygia guttata*), fire skink (*Lepidothyris fernandi*), African clawed frog, mouse, and rat. GFP-EndoN_DM_ (**Fig. 2**) and mAb735 (**Fig. 3**) positive labeling patterns were comparable, and resembled the labeling patterns expected from Müller glia. PolySia expression in adult fishes was significantly different than all other vertebrates examined, and slightly different from each other. PolySia expression in killifish was limited to the ONL, the OPL, the INL, and the S1 sublaminae of the IPL; there was also some low intensity signal in the NFL (**Fig. 2A**, **Fig. 3A**). Expression of polySia in zebrafish was more confined, and signal was observed only in the ONL, the OPL, and the very top of the INL (**Fig. 2B**, **Fig. 3B**). In adult frogs, polySia expression extended from the ONL to the NFL. PolySia in the INL exhibited a stratified pattern with arborisations in the S1, S3, and S5 IPL sublamina (**Fig. 2C**, **Fig. 3C**). In frogs, polySia signal intensity was brightest in the NFL for GFP-EndoN_DM_ labeling (**Fig. 2C**), whereas the ONL labeling was brightest when using mAb735 (**Fig. 3C**), though the overall expression patterns were the same. For the fire skink, polySia expression was strongest in the OPL and the NFL, and IPL sublaminae appeared to be thicker than those observed in frog, mice, and rats (**Fig. 2D**, **Fig. 3D)**. There was greater background fluorescence observed in fire skink retinas labeled with mAb735, compared to GFP-EndoN_DM_. PolySia expression in birds was similar to frogs and lizards. GFP-EndoN_DM_ and mAb735 labeled the ONL, OPL, IPL, and NFL, as well as S1, S3, and S5 IPL sublaminae (**Fig. 2E-F**, **Fig. 3E-F**). Background fluorescence and NFL signal was more intense in retinas labeled with mAb735 compared to GFP-EndoN_DM._ Finally, rodents had similar polySia expression patterns to frogs, lizards, and birds (**Fig. 2G-H**, **Fig 3. G-H**). Whether labeled with GFP-EndoN_DM_ or mAb735, polySia expression extended from the ONL, throughout the retina, to the NFL. A notable difference between rodents and other animals was the presence of polySia labeling in the apical processes of the Müller glia, which extend beyond the ELM into the OS space. IPL S1, S3, and S5 sublamina were also especially apparent in rodents compared to lizards and birds.

**Figure 2.**
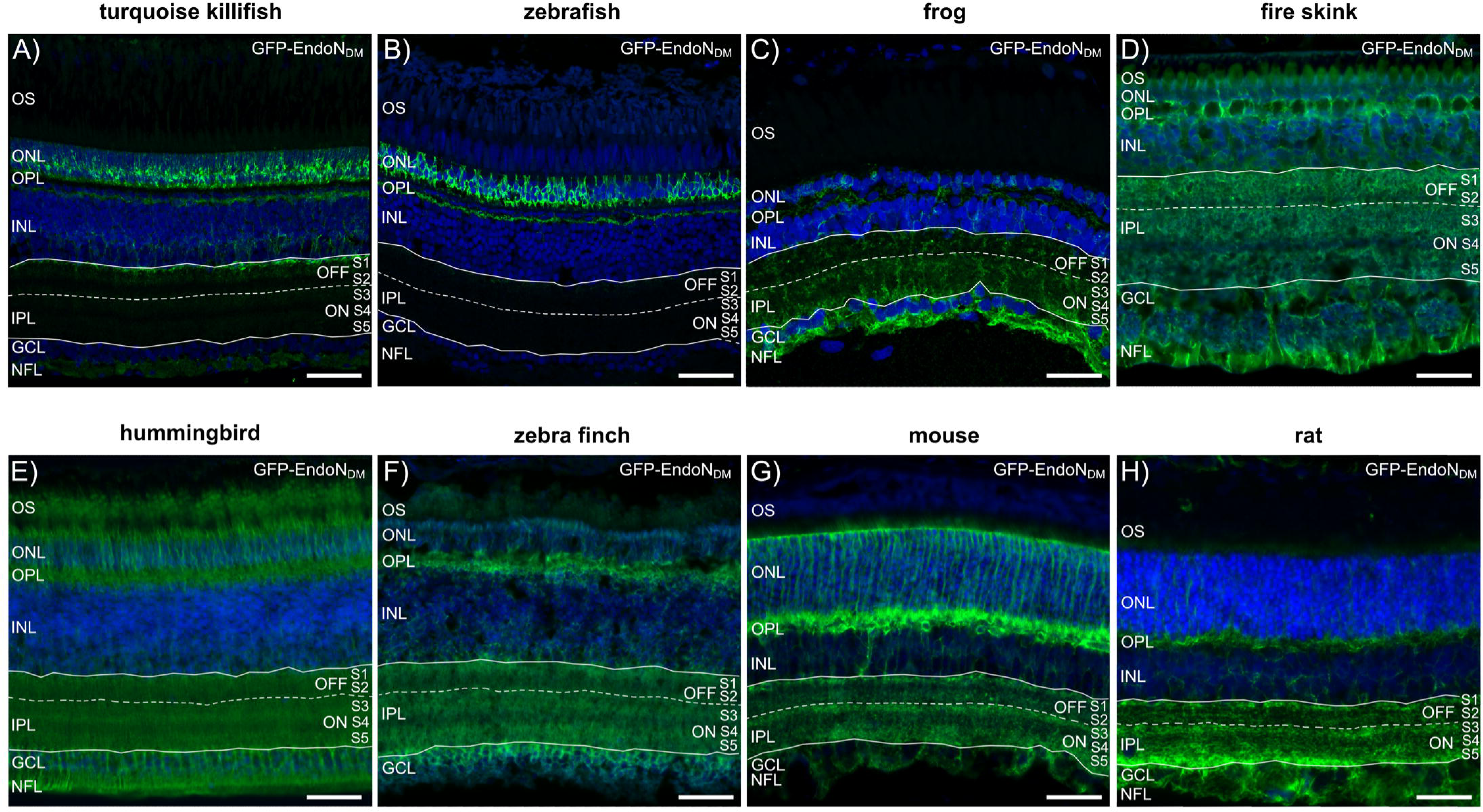
GFP-EndoN_DM_ reactivity in adult killifish (A), zebrafish (B), frog (C), fire skink (D), hummingbird (E), zebra finch (F), mouse (G), and rat (H). GFP-EndoN_DM_ expression in adult killifish or zebrafish is most concentrated to the outer retina (ONL, OPL; A,B). In the killifish, the GFP-EndoN_DM_ labeling extends to OFF sublamina S1 (A). All other species examined showed GFP-EndoN_DM_ expression throughout the entirely of the retina, likely Müller glia, from ELM to endfeet (C-H). GFP-EndoN_DM_ also exhibited increased intensity in bands in the IPL that corresponded with the S1 (OFF), S3 (ON), and S5 (ON) IPL sublaminae (C-H). *Labels*: Green, GFP-EndoN_DM_; Blue, Hoechst. *Abbreviations*: OS, outer segment layer; ONL, outer nuclear layer; OPL, outer plexiform layer; INL, inner nuclear layer; IPL, inner plexiform layer; GCL, ganglion cell layer; NFL, nerve fibre layer. Scale bar = 50 µm. n = 3-4 except fire skink (n = 1) and zebra finch (n = 2).

**Figure 3.**
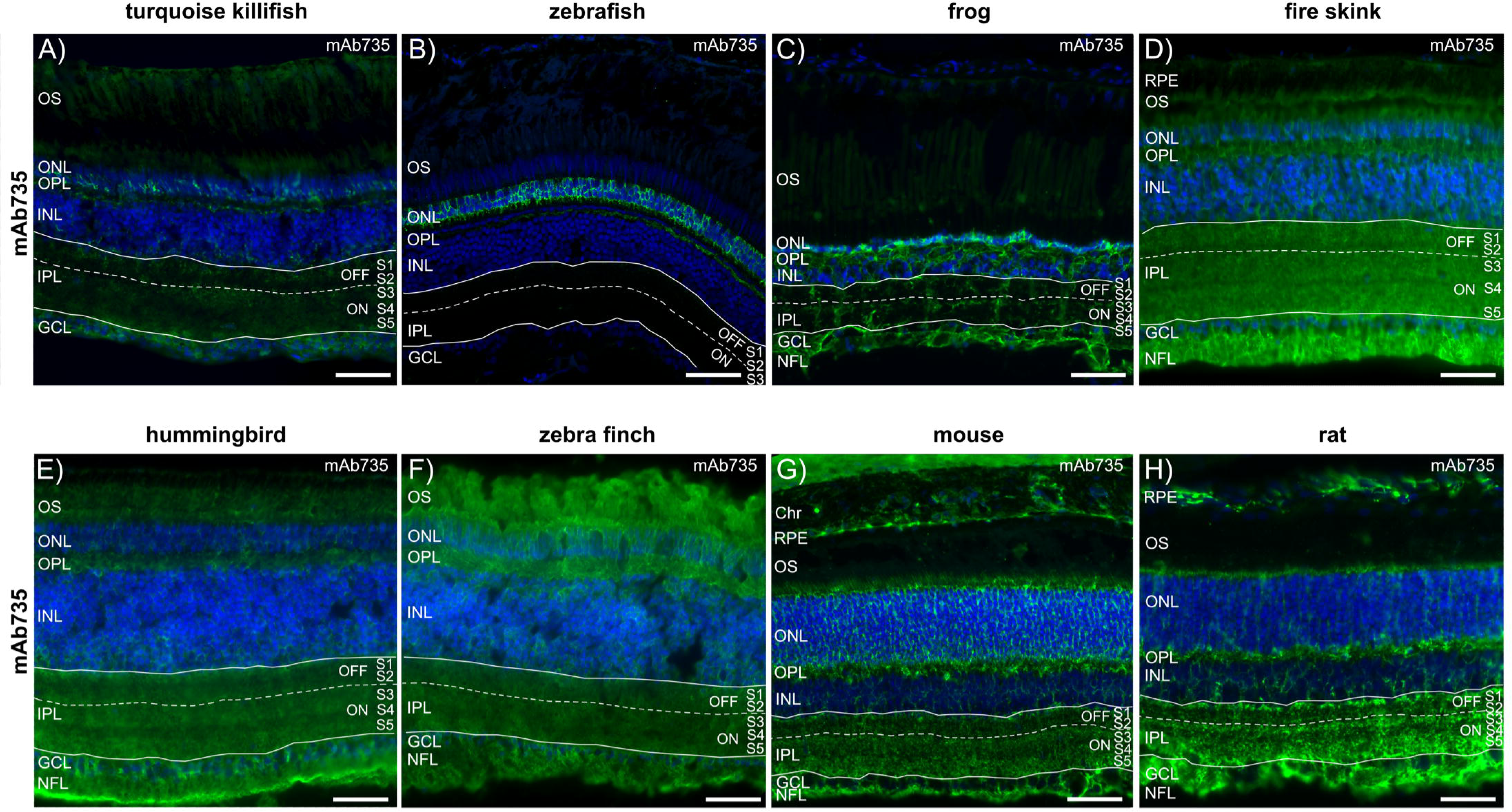
mAb735 reactivity in adult killifish (A), zebrafish (B), frog (C), fire skink (D), hummingbird (E), zebra finch (F), mouse (G), and rat (H). The labeling pattern of mAb735 in each species was the same as observed using GFP-EndoN_DM_, including restriction of signal to the outer retina in killifish and zebrafish, and labeling of the IPL S1, S3, and S5 sublaminae in all other species. mAb735 was less effective than GFP-EndoN_DM_ in highly fixed tissues such as the fire skink (D) and zebra finch (F) because background fluorescence was much higher. mAb735 also appeared to have greater non-specific labeling of photoreceptor outer segments in killifish (A), frog (C), and zebra finch (F). There was also non-specific cross reactivity of the secondary antibody (AF488 anti-mouse) in mouse and rat sclera, RPE, and intraretinal blood vessels (G-H). *Labels*: Green, mAb735; Blue, Hoechst. *Abbreviations*: Chr, choroid; RPE, retinal pigment epithelium; OS, outer segment layer; ONL, outer nuclear layer; OPL, outer plexiform layer; INL, inner nuclear layer; IPL, inner plexiform layer; GCL, ganglion cell layer; NFL, nerve fibre layer. Scale bar = 50 µm. n = 3-4 except fire skink (n = 1) and zebra finch (n = 2).

The labeling patterns of GFP-EndoN_DM_ and mAb735 in adult retinas fixed less than 24 hrs were equivalent, but mAb735 was less effective at labeling tissues that had been fixed for significant periods of time, particularly the fire skink and the zebra finch, which had high background fluorescence and less definitive Müller glia-like labeling patterns (**Fig. 3D,F**). mAb735 also appeared to have greater non-specific labeling of photoreceptor outer segments in killifish (**Fig. 3A**), frog (**Fig. 3C**), and zebra finch (**Fig. 3F**). Other slight variations in signal localization were observed between GFP-EndoN_DM_ and mAb735 due to cross-reactivity of the secondary antibody with rodent sclera, RPE, and intraretinal blood vessels (**Fig. 2G-H**, **Fig. 3G-H, Supplementary Fig. S2E**).

In addition to vertebrate retinas, we also had the opportunity to examine potential polySia expression in the retina and optic lobe of two different cephalopods: *Octopus bimaculoides* and squid (*E. berryi*). Unlike vertebrate retinas and brain, there was no significant labeling of cephalopod retinas or optic lobe using GFP-EndoN_DM_ (**Supplementary Fig. S3**).

### 3.3 GFP-EndoN_DM_ co-labeling with NCAM and established Müller glia markers

NCAM is the most common carrier of polySia in the CNS and Müller glia are known to express NCAM in zebrafish^63,86^, so we performed co-labeling with NCAM and GFP-EndoN_DM_ in adult frog to determine whether polySia expression in the retina was confined to the Müller glia. Co-labeling of NCAM and GFP-EndoN_DM_ demonstrated strong co-localization within the processes in the outer nuclear layer, cell membranes in the inner nuclear layer, and in the nerve fibre layer (**Fig. 4A-C**). GFP-EndoN_DM_ labeling was higher intensity in the outer nuclear layer compared to the rest of the retina and compared to NCAM labeling, which was more consistent across the whole retina. We also investigated the co-localization of GFP-EndoN_DM_ with vimentin in adult frog. Vimentin and GFAP label just the Müller glia cytoskeleton (**Fig. 4D-E**). Vimentin labeling co-localized with GFP-EndoN_DM_ in the inner plexiform layer, where the GFP-EndoN_DM_ signal appeared to associated with the vimentin cytoskeleton (**Fig. 4F**).

**Figure 4.**
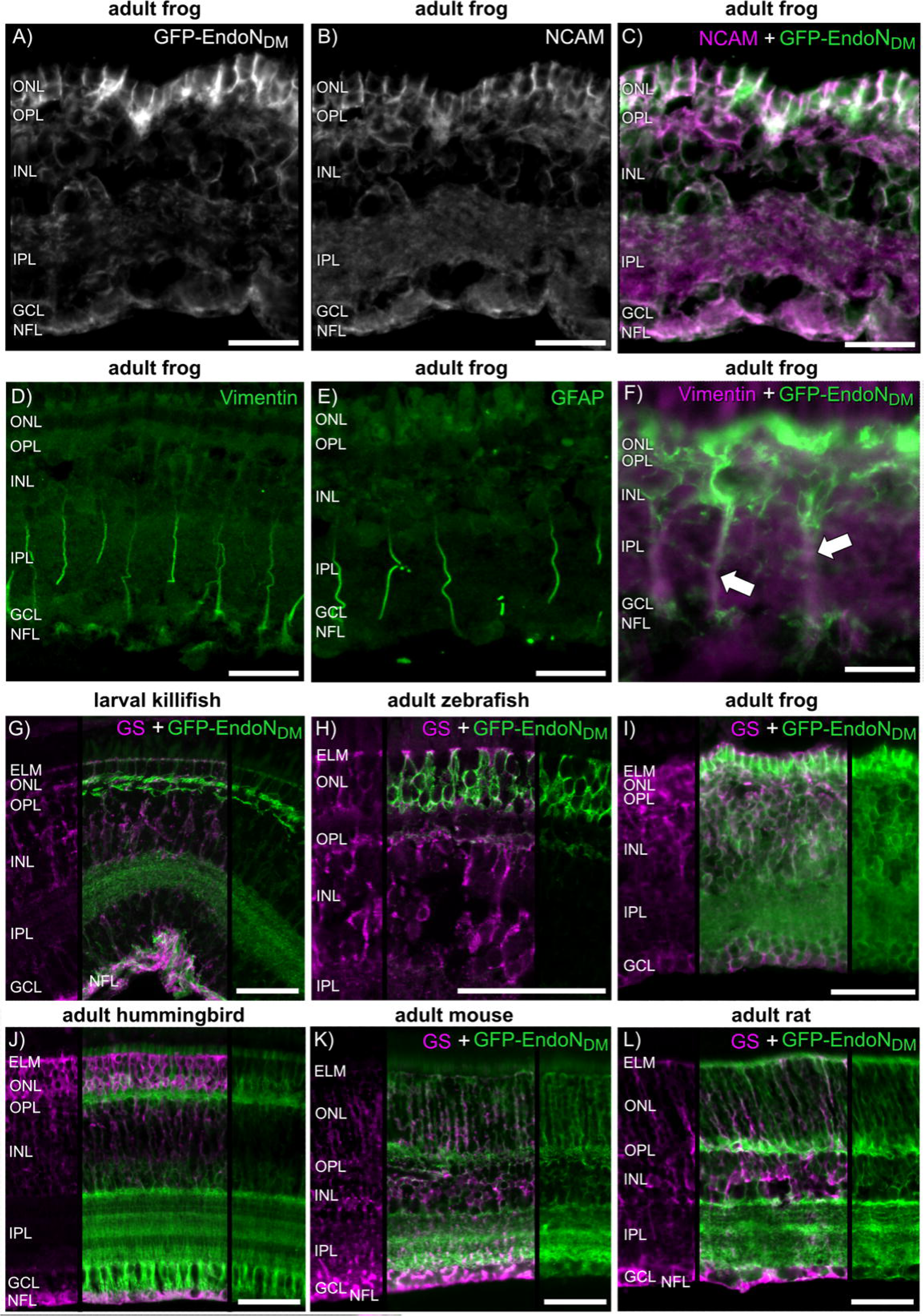
Co-labeling experiments with GFP-EndoN_DM_ and various other Müller glia markers. GFP-EndoN_DM_ co-localized with NCAM (C) and vimentin (F) in frogs, and GS in larval killifish (G), adult zebrafish (H), adult frog (I), adult hummingbird (J), adult mouse (K), and adult rat (L). There were some differences in signal intensity between GFP-EndoN_DM_ and GS. Generally, the ONL, and NFL were more intensely labeled with GS, and the OPL, IPL, and apical processes in mice and rats were more intensely labeled with GFP-EndoN_DM_. GS and GFP-EndoN_DM_ signals localized to the same spaces within the retina, except the apical processes (GFP-EndoN_DM_ in mice and rats only). *Labels*: Green, GFP-EndoN_DM_ (C,F, G-L), vimentin (D), GFAP (E); Magenta, NCAM (C), vimentin (F), GS (G-L). *Abbreviations*: GS, glutamine synthetase; ELM, external limiting membrane; ONL, outer nuclear layer; OPL, outer plexiform layer; INL, inner nuclear layer; IPL, inner plexiform layer; GCL, ganglion cell layer; NFL, nerve fibre layer. Scale bar = 50 µm. n = 3.

Finally, we performed double-labeling with GS and GFP-Endo-N_DM_ in larval killifish, adult zebrafish, adult frog, adult hummingbird, adult mouse, and adult rat. This was done to verify that polySia co-localizes with a known Müller glia marker in multiple species. GS labeled the tips of the Müller glia at the ELM, Müller glial extensions in the ONL and INL, and their endfeet with high intensity in all species examined (**Fig. 4G-L**). GS labeling of the ONL and ELM was particularly intense in the hummingbird (**Fig. 4J**). GFP-EndoN_DM_ labeling co-localized with GS at the ELM, ONL, INL, and NFL. The intensity of GS was higher in the ONL and the NFL than GFP-EndoN_DM_. There was also GFP-EndoN_DM_ signal in the apical processes of mouse and rat Müller glia (**Fig. 4K-L**). The most apparent co-localization of GS and GFP-EndoN_DM_ occurred in the ELM, ONL, INL, and NFL. There was less co-localization of GS and GFP-EndoN_DM_ in the OPL and the IPL, where GFP-EndoN_DM_ signal was more intense and demonstrated IPL sublaminae organization where GS did not exhibit significant IPL sublaminae signal. We did not perform double-labeling with mAb735 because all primary antibodies were mouse monoclonal antibodies and the labeling patterns for GFP-EndoN_DM_ and mAb735 were effectively indistinguishable.

### 3.4 GFP-EndoN_DM_ labeling in a mouse model of severe retinal degeneration

To test whether polySia labeling of Müller glia could also visualize gliosis, we obtained eyes from aryl hydrocarbon receptor-interacting protein-like 1 knockout (*Aipl1^-/-^*) mice, aged P97. These mice are a model for human Leber Congential Amaurosis (LCA), and have been reported previously to exhibit rapid photoreceptor degeneration, loss of electroretinogram (ERG), and signs of Müller gliotic changes, as early as P14^87^. We were able to visualize gliotic changes in these mice using GFP-EndoN_DM_ and mAb735 (**Fig. 5**). Double-labeling with GS showed strong correlation between high intensity labeling patterns near the outer plexiform layer and in the endfeet. As seen with healthy tissues, GFP-EndoN_DM_ had increased signal around cell bodies in the inner nuclear layer and the IPL sublaminae labeling remained intact (**Fig. 5A-B,D**).

**Figure 5.**
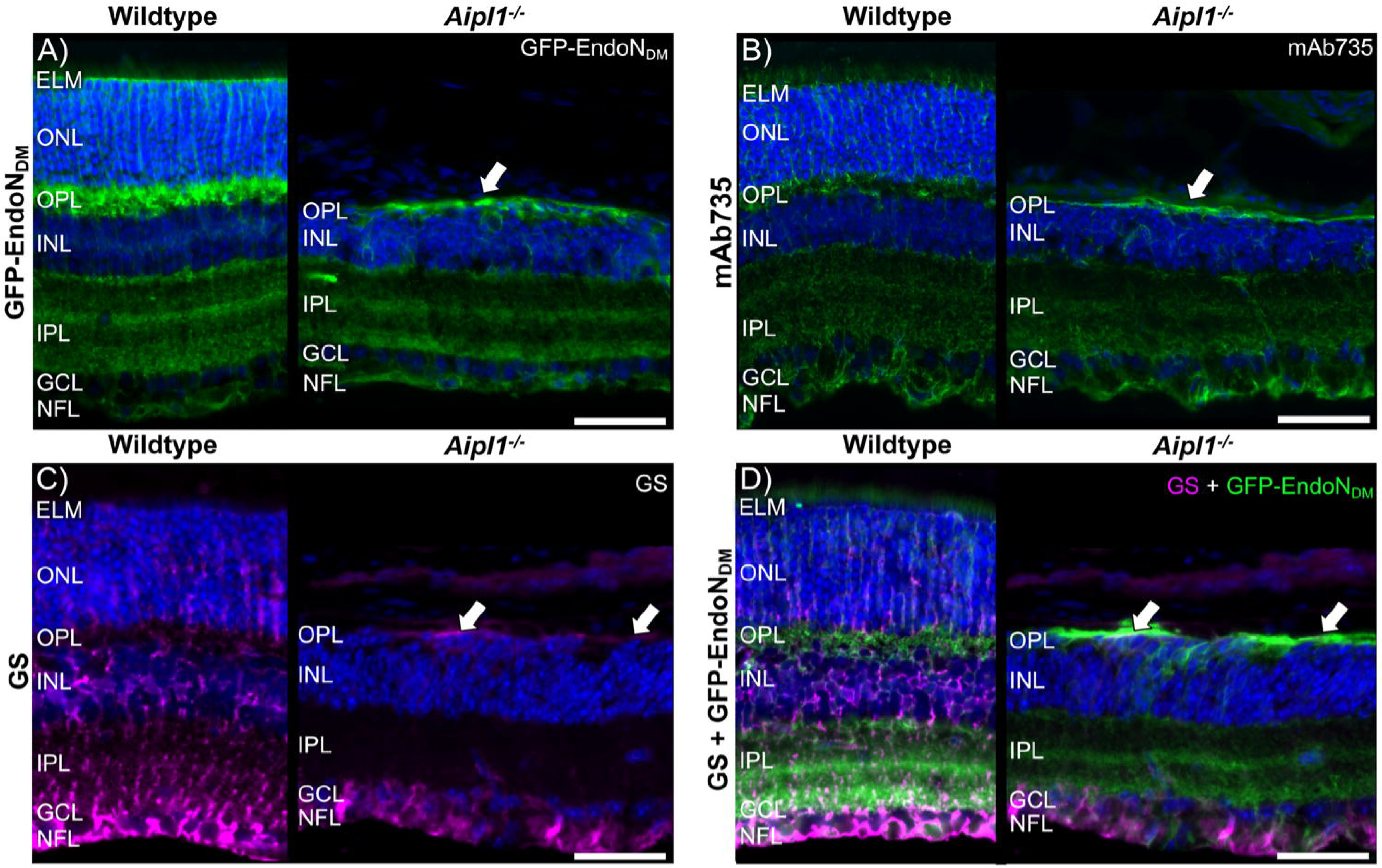
Detection of gliosis in a 97 day old *Aipl1^-/-^* mouse by GFP-EndoN_DM_ (A), mAb735 (B), glutamine synthetase (GS; C), and double-labeling with GFP-EndoN_DM_ + GS (D). *Aipl1^-/-^* mice had complete loss of their outer segment and outer nuclear layer, and gliotic scarring in the OPL was visible using all markers (white arrows). GS and GFP-EndoN_DM_ labeling of scarring was co-localized (D). *Labels*: Green, GFP-EndoN_DM_ (A,D), mAb735 (B); Magenta, GS; Blue, Hoechst. *Abbreviations*: GS, glutamine synthetase; ELM, external limiting membrane; ONL, outer nuclear layer; OPL, outer plexiform layer; INL, inner nuclear layer; IPL, inner plexiform layer; GCL, ganglion cell layer; NFL, nerve fibre layer. Scale bar = 50 µm. n = 3.

### 3.5 GFP-EndoN_DM_ localization in aged human retina and retinal organoids

Finally, we tested GFP-EndoN_DM_ localization in human retina and patient-derived retinal organoids; mAb735 was not tested due to tissue scarcity. The labeling pattern of GFP-EndoN_DM_ is very similar to what is seen in other vertebrate retinas, including the IPL sublaminae (**Fig. 6A-C**). The bright band of labeling at the ELM was not as distinct as observed in other vertebrates, but was most visible in tissues sampled from just outside of the macula (**Fig. 6B**). One sample also exhibited choroidal Schwann cells that labeled intensely with GFP-EndoN_DM_ (**Fig. 6A, D-E**). There was significant red and green autofluorescence in the RPE (**Fig. 6A-E).**

**Figure 6.**
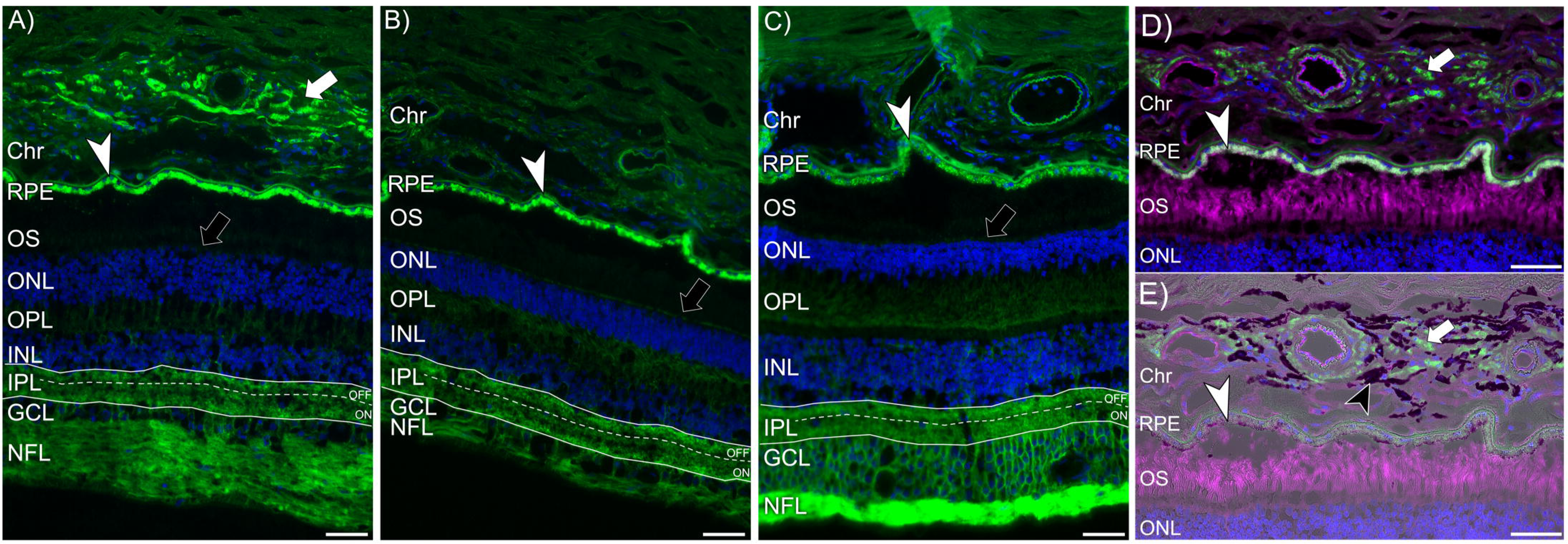
GFP-EndoN_DM_ labeling in post-mortem human retina sections that are outside of the macula (A), close to the macula (B), and close to the optic nerve (C). The labeling pattern is very similar to what is seen in other vertebrate retinas, including the IPL sublaminae, but the bright band of labeling at the ELM is not as distinct as observed in other vertebrates in sections taken close to the macula and the optic nerve. Choroidal Schwann cells labeled brightly with GFP-EndoN_DM_ in one of the samples (A, D-E, white arrow), and did not co-localize with melanocytes (E, black cells, black arrowhead). There was significant red and green channel RPE autofluorescence observed in all samples (A-E, white arrowheads). Labels: Magenta, wheat germ agglutinin (WGA); Green, GFP-EndoN_DM_; Blue, Hoechst. *Abbreviations*: Chr, choroid; RPE, retinal pigment epithelium; OS, outer segment layer; ONL, outer nuclear layer; OPL, outer plexiform layer; INL, inner nuclear layer; IPL, inner plexiform layer; GCL, ganglion cell layer; NFL, nerve fibre layer. Scale bar = 50 µm. n = 3.

Wildtype and Parkinson’s patient-derived retinal organoids also exhibited strong expression of GFP-EndoN_DM_. Labeling was dispersed throughout the organoid in intensely-labelled areas that resembled plaques, laminations, and the external limiting membrane (**Fig. 7**). ELM-like structures were especially apparent along the outer organoid surface, where photoreceptor outer segments should be, and near photoreceptor rosettes that were present inside of the organoids. It also appeared as if more plaques were present in patient-derived organoids compared to healthy ones, although this was not quantified.

**Figure 7.**
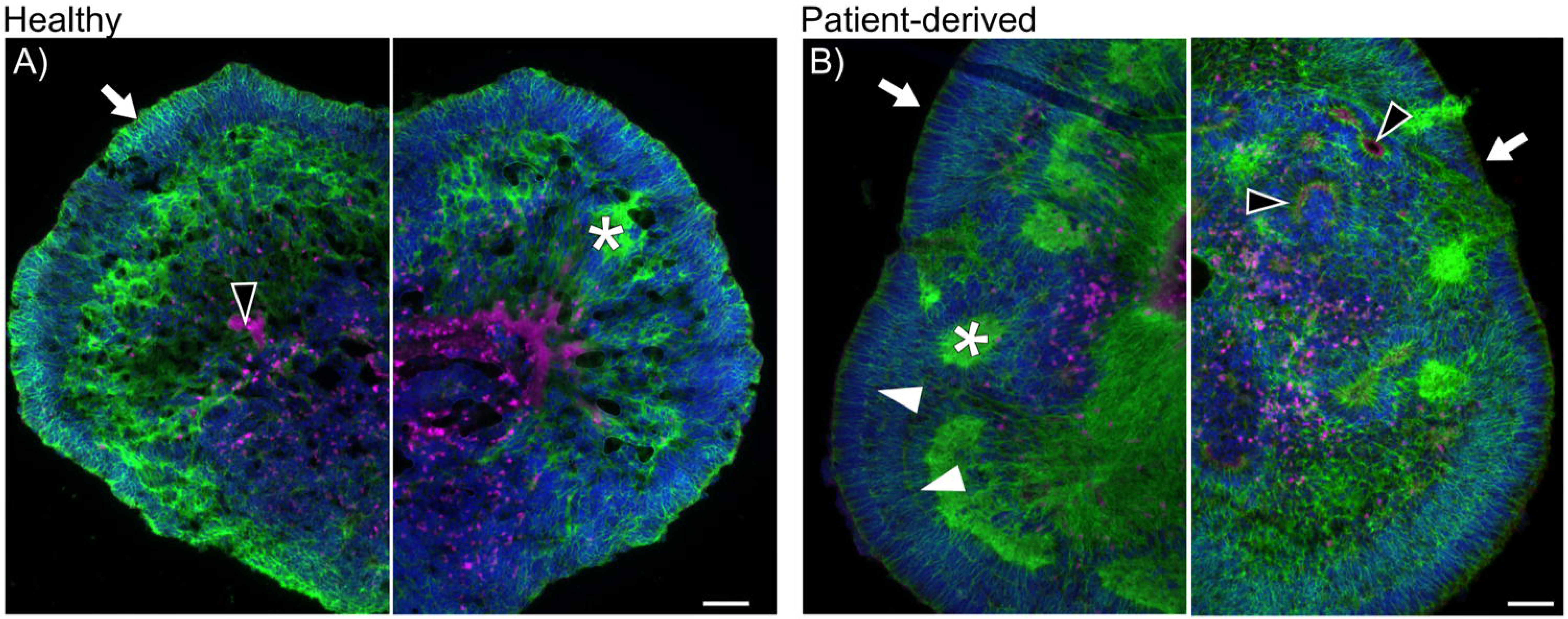
GFP-EndoN_DM_ labeling in cross sections of healthy (A) and patient-derived (B) retinal organoids, aged 120 days. Expression patterns resembled external limiting membrane-like labeling (white arrows), plaque-like areas of high intensity labeling (asterisks), and areas of high intensity surrounding rosettes (black arrowheads). *Labels*: Magenta, wheat germ agglutinin (WGA); Green, GFP-EndoN_DM_; Blue, Hoechst. Scale bar = 50 µm. n = 3

## 4. Discussion

### 4.1 PolySia during development and homeostasis across vertebrate species

In this study, we showed that GFP-EndoN_DM_ and mAb735, which label polySia, are effective Müller glia markers in most common model organisms, retinal organoids, and human tissues. We report that polySia is expressed throughout the entirely of the Müller glia cell body, from ELM to endfeet, with fine banding in the S1, S3, and S5 IPL sublaminae in healthy retinas of humans and non-fish vertebrates. Mice and rats also exhibit significant polySia expression in the Müller glia apical processes that extend beyond the ELM. Interestingly, polySia labeling in zebrafish and killifish is restricted to a very defined region of the glia in the outer retina. The reason for this restriction of expression compared to other vertebrates remains unknown.

Our data agree with previous reports of polySia-NCAM expression in the retina of adult zebrafish^63^, salamander (*Pleurodeles waltl*)^62^, frog (*Lithobates pipiens*)^61^, chicken^88^, and rats^60^. These data are, to our knowledge, the first reports of polySia expression in larval and adult killifish retinas. We found that α2,8-Neu5Ac polySia is not significantly expressed in cephalopod retinas or optic lobes. There is little literature on sialic acids in mollusks, as it is generally accepted that they do not express them. However, one previous report indicated the presence of Neu5Ac in *Octopus vulgaris* and *Todarodes pacificus* hepatopancreatic and cerebral ganglia tissues, on the basis of negative ion electrospray ionization mass spectrometry (ESI-MS) and fluorometric high-performance liquid chromatography (HPLC)^89^.

We believe that retinal polySia detected by GFP-EndoN_DM_ and mAB735 corresponds primarily to Müller glia on the basis of co-labeling experiments with NCAM, vimentin, and GS, which demonstrated significant overlap with GFP-EndoN_DM_. It should be noted, however, that although the predominant form of CNS-associated polySia is polySia-NCAM, other retinal proteins with polySia modification include an alpha subunit of voltage-gated sodium channels, CD36, synaptic cell adhesion molecule 1 (SynCAM 1), and neuropillin-2^90–93^. SynCAM 1 and voltage-gated sodium channels are particularly abundant in the retina, and could contribute to polySia labeling near the photoreceptor nuclei and the ganglion cell axons^94–97^. Retinal SynCAM 1 does not appear to co-localize with Müller glia or GFAP, however^94,95^.

While the presence of polySia-NCAM in the brain during development and adulthood has been well-studied^44,52,53,58^, there are comparatively fewer studies on changes in polySia-NCAM expression in the retina over time or with disease. Most studies in the brain report that polySia levels are high during embryonic development when cell migration and plasticity is most active, and then levels decrease as animals age, with the exception of some areas where plasticity and neurogenesis remain important^54–58,98^. These developmental changes are attributed to roles of polySia in cell migration^99^, axon growth, guidance and fasciculation^100–102^, synaptic plasticity^103^, and dendritic sprouting^104,105^. Most of the animals assessed in our report were adults; however, we were able to compare retinal polySia labeling between early developmental and adult life stages for killifish, zebrafish, frogs, mice, and rats. PolySia was present in Müller glia during early retinal development, before the retina is fully laminated, and in adulthood. In most vertebrate species examined, the intensity of polySia labeling within the retina did not change significantly between larval/post-natal and adult life stages. The exception to this is the bony fishes. PolySia was absent in free-swimming larval zebrafish and restricted to the outer retina in adults. On the other hand, polySia expression was more widespread in the larval killifish retina including the ONL, OPL, INL, IPL, and NFL, compared to the adult killifish retina where it was restricted to the outer retina and the S1 IPL sublamina. We observed polySia labelling in the larval zebrafish, killifish, and frog brains in agreement with previous reports of high polySia expression in the vertebrate brain during embryonic development^44,52,53,58^. Zebrafish are popular aquatic model organisms for the study of retinal development and disease^106^, and interest in killifish as a model of retinal and brain disease is increasing due to their unique lifecycle and fast aging^107–109^, so the finding that polySia expression in the Müller glia of fishes differs from other vertebrates, even amphibians, is a noteworthy consideration for future studies using these animals.

The IPL is divided into ON-(S3, S4, and S5) and OFF-(S1 and S2) sublaminae, which represent termination layers of rod and cone bipolar-specific circuitry. Here, we report polySia labeling of fine arborisations in the S1, S3, S5 IPL sublaminae in frog, lizard, birds, rodents, and humans. Our data align with previous reports of Müller glia fine processes in the IPL sublaminae in a number of other species including cat, dog, echidna, guinea pig, rabbit, pigeon, chicken, and monkey^88,110–112^. Recently, a transgenic mouse was used to confirm that the Müller glia-associated IPL sublaminae are located above, between, and below the ON and OFF cholinergic acetyltransferase (ChAT) positive amacrine cell layers^32^, in the S1 OFF-sublamina and the S3 and S5 ON-sublaminae^113,114^. Although fish also have these IPL sublaminae, and S1 was labeled with GFP-EndoN_DM_ in killifish, S3 and S5 were not significantly labeled using polySia in either fish species examined. This is likely a difference in protein modification with polySia between fishes and other vertebrates, rather than significant differences in retinal circuitry between bony fishes and other vertebrates.

In human tissues, the labeling of the ELM was not as apparent as other vertebrates examined. This could be due to location within the eye, as tissue sections from just outside of the macula had the best labeling of the ELM. It could also be due to age, since these tissue sections represented aged humans (69-81 years), in comparison to the early to mid adulthood tissue sections that were taken from all other vertebrates examined. There was also significant red-green autofluorescence in the RPE of the human retinas, which is typical of aged retinas due to the build-up of lipofuscins and other indigestible protein and lipid waste products^115,116^. One human retina also demonstrated additional labeling in a subset of choroidal cells that was not seen in other species. We suspect that these were Schwann cells because they did not co-localize with choroidal melanocytes, and non-myelinating Schwann cells are present in the choroid and have been reported previously to express polySia-NCAM^46,117^. Schwann cells are a type of glial cell that is not usually found in healthy retinas, but instead are usually associated with peripheral nervous system development and repair^118^. They can infiltrate into the retina under conditions of photoreceptor degeneration and high oxidative stress, however, and injection of Schwann cells has been shown to be protective against retinal ganglion cell axon degeneration in Royal College of Surgeons rats^119,120^.

#### PolySia in disease states

We report here, that polySia labeling is also useful for studying gliosis, which is a reactive response observed in Müller glia and astrocytes to various forms of nervous system injury and disease^22,121,122^. In cases of severe reactive gliosis, Müller cells lose functionality and form glial scars, inhibiting neuronal regeneration and survival. These gliotic scars were detected using GFP-EndoN_DM_ in *Aipl1^-/-^* mice, and were comparable to previously published results in the same mouse model using GFAP and GS to detect gliosis^87^.

Finally, we showed that polySia is an effective marker for Müller glia-like cells in retinal organoids derived from human cell lines, and that the labeling patterns of polySia in retinal organoids closely resembled those of vimentin^72^. A recent study investigated the impacts of the loss of Neu5Ac on cortical organoid development and found that organoids with mutations in *N*-acetylneuraminic acid synthase, an enzyme critical for the synthesis of polySia, had significantly reduced proliferation, organization, and synaptic connectivity^123^. Although the role of polySia in developing and adult Müller glia is still unknown, a number of studies have shown that intravitreal injections of polySia are neuroprotective under conditions of AMD-like disease and kainic acid-induced retinal ganglion cell excitotoxity^124–126^. It is possible that the Müller glia may mediate this effect.

#### Antibodies vs. lectins

Because Müller glia play such a critical role in retinal physiology, it is important to have effective tools to study them. Common labeling tools, such as primary antibodies against GFAP, vimentin, and GS can sometimes have limitations in their applicability across various animal models. First, there can be significant differences in protein structure between species, which can affect the ability of antibodies to bind to their target. Most commercially available antibodies are made to cross-react with human, mouse, and rat proteins, because they are the most common animal and cell model organisms. While effective in these models, commercial antibodies rarely adequately label protein targets from other species, such as frogs, fish, or birds. The structure of polySia is highly conserved across vertebrates, eliminating these issues with poor protein structure conservation across species.

Second, some antigens are poorly immunogenic, of limited availability, or are unstable, and raising antibodies against them is therefore difficult. For example, *Xenopus laevis* may not express GFAP, and instead expresses a protein that cross-reacts with both GFAP and vimentin antibodies^127^. Furthermore, antibodies for unique model organisms are difficult, time consuming, and expensive to try to generate, particularly if the sequence of the proteins of interest is unknown or poorly annotated. For instance, the killifish genome currently lacks a curated database like those available for most other model organisms such as https://Xenbase.org, https://ZFIN.org, https://Flybase.org, and the mouse genome database (MGD), which makes designing custom antibodies difficult.

Third, the molecular targets of GFAP, vimentin, and GS may not be ideal for visualization of whole Müller glia cell bodies and fine processes. GFAP and vimentin are cytoskeletal markers, and therefore do not label cell membranes or Müller glial arborisations. In fact, GFAP labels only ∼15% of the volume of astrocytes, and has similar efficacy for Müller glia^31^. GS is a cytoplasmic marker and is therefore not detected in Müller glia cell membranes or fine IPL arborisations, resulting in missing morphological information. Labeling of GS can also be dependent on the upregulation of the target protein glutamine synthetase, such as during disease, for best visualization^29,128,129^. Nevertheless, we consider GS to be the better antibody for Müller glia visualization. The GS polyclonal antibody used here cross-reacts with many different species, and it better highlights Müller glial morphology than either vimentin or GFAP antibodies.

As for the polySia markers tested here, we consider GFP-EndoN_DM_ to be a better marker than mAb735. GFP-EndoN_DM_ is a conjugate, therefore does not have cross-reactivity problems due to non-specific binding of the mouse secondary antibody to endogenous IgGs. It can also be used in double-labeling experiments with any species of primary antibody or other lectin, leading to greater versatility in experimental design. Further advantages to GFP-EndoN_DM_ are that it is inexpensive to synthesize in large quantities, far more stable than antibodies, resistant to heat treatment and detergents, and can be engineered to have different conjugates. For example, the GFP can be switched out for a biotinylated version of EndoN_DM_ that could be coupled to any fluorescent streptavidin or streptavidin-coated agarose beads if one wanted to enrich Müller glia from a dissociated retina sample^79^. Finally, binding of GFP-EndoN_DM_ to polySia appears to be resistant to formaldehyde over-fixation, as evidenced by its successful use in our study on tissues that have been fixed for a significant amount of time (lizard and zebra finch) compared to poorer mAb735 labeling of the same tissues. Direct conjugates can sometimes have low signal due to 1:1 binding with the target, unlike the signal amplification from combined use of primary and secondary antibodies. However, because GFP-EndoN_DM_ binds to the α2,8 linkages of polySia chains, it has the opportunity to bind tens to hundreds of times along these chains, resulting in significant signal amplification similar to primary-secondary systems.

## Conclusion

Here, we report the versatility of polySia as a marker of Müller glia in amphibian, avian, mammalian, human, and retinal organoid tissues. Labeling polySia is convenient, simple, and useful for the study of Müller glia across different species, developmental stages, and disease states. We suggest GFP-EndoN_DM_ and mAb735 as suitable alternatives or supplemental markers to conventional Müller glia markers such as GFAP, vimentin, and GS in models or experimental paradigms where using these reagents is inconvenient or not possible.

## Acknowledgements

We thank the donors to the Iowa Lions Eye bank for their generosity and commitment to science, for they have given the incredible gift of a part of themselves so that we may learn more about the human body. Thank you to the following persons for providing human eyes, animal eyes, and retinal organoids, without which this project would not have been possible: Maryam Hejazi (University of Alberta; adult wildtype mice), Dr. Daniel Hass (University of Washington, Hurley Lab; *Aipl1^-/-;-/+^* mice), Dr. Christine Webber (University of Alberta; adult rat), Nichole Vestby (University of Alberta, Director of Operations NCAS), Dr. Matthew Macauley (University of Alberta; postnatal mouse), Dr. Francis Plane (University of Alberta; postnatal rat), Dr. Zach Hall (in memoriam; University of Alberta; larval zebrafish), Dr. Cristian Gutierrez (University of Alberta, Wylie Lab; hummingbird, fire skink), Dr. Judit Pungor (University of Oregon, Niell Lab; cephalopods), Dr. Sandra Tenreiro (NOVA Medical School, Universidade Nova de Lisboa; retinal organoids), and Dr. Robert F. Mullins (University of Iowa; human retinas). We also wish to acknowledge the Marine Resources Center (Woods Hole, MA, USA) for the husbandry and care of cephalopods for the greater research community.

## References

1. Reichenbach, A. & Bringmann, A. New functions of Müller cells. Glia 61, 651–678 (2013).

2. Xue, Y. et al. The role of retinol dehydrogenase 10 in the cone visual cycle. Sci. Rep. 7, 2390 (2017).

3. Poitry-Yamate, C., Poitry, S. & Tsacopoulos, M. Lactate released by Muller glial cells is metabolized by photoreceptors from mammalian retina. J. Neurosci. 15, 5179–5191 (1995).

4. Tworig, J. M. & Feller, M. B. Müller Glia in Retinal Development: From Specification to Circuit Integration. Front. Neural Circuits 15, 815923 (2022).

5. Sakami, S., Imanishi, Y. & Palczewski, K. Müller glia phagocytose dead photoreceptor cells in a mouse model of retinal degenerative disease. FASEB J. 33, 3680–3692 (2019).

6. Bassett, E. A. & Wallace, V. A. Cell fate determination in the vertebrate retina. Trends Neurosci. 35, 565–573 (2012).

7. Reichenbach, A. et al. What do retinal Müller (glial) cells do for their neuronal ‘small siblings’? J. Chem. Neuroanat. 6, 201–213 (1993).

8. Newman, E. & Reichenbach, A. The Müller cell: a functional element of the retina. Trends Neurosci. 19, 307–312 (1996).

9. Dubois-Dauphin, M. et al. Early postnatal Müller cell death leads to retinal but not optic nerve degeneration in NSE-Hu-Bcl-2 transgenic mice. Neuroscience 95, 9–21 (1999).

10. Byrne, L. C., et al. AAV-Mediated, Optogenetic Ablation of Müller Glia Leads to Structural and Functional Changes in the Mouse Retina. PLoS ONE 8, e76075 (2013).

11. Shen, W. et al. Conditional Müller Cell Ablation Causes Independent Neuronal and Vascular Pathologies in a Novel Transgenic Model. J. Neurosci. 32, 15715–15727 (2012).

12. Nilsson, S. E. RECEPTOR CELL OUTER SEGMENT DEVELOPMENT AND ULTRASTRUCTURE OF THE DISK MEMBRANES IN THE RETINA OF THE TADPOLE (RANA PIPIENS). J Ultrastruct Res 11, 581–602 (1964).

13. Bringmann, A. & Wiedemann, P. Müller Glial Cells in Retinal Disease. Ophthalmologica 227, 1–19 (2012).

14. Pfeiffer, R. L. et al. Pathoconnectome Analysis of Müller Cells in Early Retinal Remodeling. Adv. Exp. Med. Biol. 1185, 365–370 (2019).

15. Reichenbach, A. et al. The structure of rabbit retinal Müller (glial) cells is adapted to the surrounding retinal layers. Anat. Embryol. (Berl.) 180, 71–79 (1989).

16. Omri, S. et al. The outer limiting membrane (OLM) revisited: clinical implications. Clin. Ophthalmol. Auckl. NZ 4, 183–195 (2010).

17. Zhang, K. Y. & Johnson, T. V. The internal limiting membrane: Roles in retinal development and implications for emerging ocular therapies. Exp. Eye Res. 206, 108545 (2021).

18. Hamon, A. et al. Linking YAP to Müller Glia Quiescence Exit in the Degenerative Retina. Cell Rep. 27, 1712–1725.e6 (2019).

19. Hamon, A., Roger, J. E., Yang, X. & Perron, M. Müller glial cell-dependent regeneration of the neural retina: An overview across vertebrate model systems. Dev. Dyn. 245, 727–738 (2016).

20. Wan, J. & Goldman, D. Retina regeneration in zebrafish. Curr. Opin. Genet. Dev. 40, 41–47 (2016).

21. Bringmann, A. et al. Cellular signaling and factors involved in Müller cell gliosis: Neuroprotective and detrimental effects. Prog. Retin. Eye Res. 28, 423–451 (2009).

22. Bringmann, A. et al. Müller cells in the healthy and diseased retina. 25, 397–424 (2006).

23. Bringmann, A. et al. Role of retinal glial cells in neurotransmitter uptake and metabolism. Neurochem. Int. 54, 143–160 (2009).

24. Navneet, S., Wilson, K. & Rohrer, B. Müller Glial Cells in the Macula: Their Activation and Cell-Cell Interactions in Age-Related Macular Degeneration. Investig. Opthalmology Vis. Sci. 65, 42 (2024).

25. Edwards, M. M. et al. Subretinal Glial Membranes in Eyes With Geographic Atrophy. Investig. Opthalmology Vis. Sci. 58, 1352 (2017).

26. Edwards, M. M. et al. Idiopathic preretinal glia in aging and age-related macular degeneration. Exp. Eye Res. 150, 44–61 (2016).

27. Palko, S. I., et al. Compartmentalized citrullination in Muller glial endfeet during retinal degeneration. Proc. Natl. Acad. Sci. 119, e2121875119 (2022).

28. Bignami, A. & Dahl, D. The radial glia of Müller in the rat retina and their response to injury. An immunofluorescence study with antibodies to the glial fibrillary acidic (GFA) protein. Exp. Eye Res. 28, 63–69 (1979).

29. Eisenfeld, A. J., Bunt-Milam, A. H. & Sarthy, P. V. Müller cell expression of glial fibrillary acidic protein after genetic and experimental photoreceptor degeneration in the rat retina. Invest. Ophthalmol. Vis. Sci. 25, 1321–1328 (1984).

30. Chen, H. & Weber, A. J. Expression of glial fibrillary acidic protein and glutamine synthetase by Müller cells after optic nerve damage and intravitreal application of brain-derived neurotrophic factor. Glia 38, 115–125 (2002).

31. Bushong, E. A., Martone, M. E., Jones, Y. Z. & Ellisman, M. H. Protoplasmic Astrocytes in CA1 Stratum Radiatum Occupy Separate Anatomical Domains. J. Neurosci. 22, 183–192 (2002).

32. Wang, J. et al. Anatomy and spatial organization of Müller glia in mouse retina. J. Comp. Neurol. 525, 1759–1777 (2017).

33. Sarthy, P. V., Fu, M. & Huang, J. Developmental expression of the Glial fibrillary acidic protein (GFAP) gene in the mouse retina. Cell. Mol. Neurobiol. 11, 623–637 (1991).

34. Stanley, P., Moremen, K. W., Lewis, N. E., Taniguchi, N. & Aebi, M. N-Glycans. in Essentials of Glycobiology (eds Varki, A., et al.) (Cold Spring Harbor Laboratory Press, Cold Spring Harbor (NY), 2022).

35. Sato, C. & Kitajima, K. Polysialylation and disease. Mol. Aspects Med. 79, 100892 (2021).

36. Marki, A., Esko, J. D., Pries, A. R. & Ley, K. Role of the endothelial surface layer in neutrophil recruitment. J. Leukoc. Biol. 98, 503–515 (2015).

37. Ito, T. et al. Recognition of N-glycolylneuraminic acid linked to galactose by the alpha2,3 linkage is associated with intestinal replication of influenza A virus in ducks. J. Virol. 74, 9300–9305 (2000).

38. Nemanichvili, N. et al. Wild and domestic animals variably display Neu5Ac and Neu5Gc sialic acids. Glycobiology (2022) doi:10.1093/glycob/cwac033.

39. Davies, L. R. L. et al. Metabolism of vertebrate amino sugars with N-glycolyl groups: resistance of α2-8-linked N-glycolylneuraminic acid to enzymatic cleavage. J. Biol. Chem. 287, 28917–28931 (2012).

40. Angata, K. & Fukuda, M. Roles of Polysialic Acid in Migration and Differentiation of Neural Stem Cells. in Methods in Enzymology vol. 479 25–36 (Elsevier, 2010).

41. Heimburg-Molinaro, J. et al. Cancer vaccines and carbohydrate epitopes. Vaccine 29, 8802–8826 (2011).

42. Hunter, C. et al. Attenuation of Polysialic Acid Biosynthesis in Cells by the Small Molecule Inhibitor 8-Keto-sialic acid. ACS Chem. Biol. 18, 41–48 (2023).

43. Finne, J. Occurrence of unique polysialosyl carbohydrate units in glycoproteins of developing brain. J. Biol. Chem. 257, 11966–11970 (1982).

44. Rothbard, J. B., Brackenbury, R., Cunningham, B. A. & Edelman, G. M. Differences in the carbohydrate structures of neural cell-adhesion molecules from adult and embryonic chicken brains. J. Biol. Chem. 257, 11064–11069 (1982).

45. Pestronk, A., Schmidt, R. E., Bucelli, R. & Sim, J. Schwann cells and myelin in human peripheral nerve: Major protein components vary with age, axon size and pathology. Neuropathol. Appl. Neurobiol. 49, e12898 (2023).

46. Roche, P.-H. et al. Expression of cell adhesion molecules in normal nerves, chronic axonal neuropathies and Schwann cell tumors. J. Neurol. Sci. 151, 127–133 (1997).

47. Edelman, G. M. & Crossin, K. L. CELL ADHESION MOLECULES: Implications for a Molecular Histology. Annu. Rev. Biochem. 60, 155–190 (1991).

48. Rutishauser, U. Polysialic acid in the plasticity of the developing and adult vertebrate nervous system. Nat. Rev. Neurosci. 9, 26–35 (2008).

49. Mindler, K., Ostertag, E. & Stehle, T. The polyfunctional polysialic acid: A structural view. Carbohydr. Res. 507, 108376 (2021).

50. Rutishauser, U. & Landmesser, L. Polysialic acid in the vertebrate nervous system: a promoter of plasticity in cell-cell interactions. Trends Neurosci. 19, 422–427 (1996).

51. Sato, C. & Kitajima, K. Disialic, oligosialic and polysialic acids: distribution, functions and related disease. J. Biochem. (Tokyo) 154, 115–136 (2013).

52. Bartsch, U., Kirchhoff, F. & Schachner, M. Highly sialylated N-CAM is expressed in adult mouse optic nerve and retina. J. Neurocytol. 19, 550–565 (1990).

53. Hildebrandt, H. et al. Polysialic acid on the neural cell adhesion molecule correlates with expression of polysialyltransferases and promotes neuroblastoma cell growth. Cancer Res. 58, 779–784 (1998).

54. Amoureux, M.-C. et al. Polysialic Acid Neural Cell Adhesion Molecule (PSA-NCAM) is an adverse prognosis factor in glioblastoma, and regulates olig2 expression in glioma cell lines. BMC Cancer 10, 91 (2010).

55. Petridis, A. K., El Maarouf, A. & Rutishauser, U. Polysialic acid regulates cell contact-dependent neuronal differentiation of progenitor cells from the subventricular zone. Dev. Dyn. 230, 675–684 (2004).

56. Rutishauser, U. & Landmesser, L. Polysialic acid in the vertebrate nervous system: a promoter of plasticity in cell-cell interactions. Trends Neurosci. 19, 422–427 (1996).

57. Durbec, P. & Cremer, H. Revisiting the function of PSA-NCAM in the nervous system. Mol. Neurobiol. 24, 53–64 (2001).

58. Yang, Y. et al. Comparative Studies of Polysialic Acids Derived from Five Different Vertebrate Brains. Int. J. Mol. Sci. 21, 8593 (2020).

59. Murphy, J. A., Hartwick, A. T. E., Rutishauser, U. & Clarke, D. B. Endogenous Polysialylated Neural Cell Adhesion Molecule Enhances the Survival of Retinal Ganglion Cells. Invest. Ophthalmol. Vis. Sci. 50, 861–869 (2009).

60. Sawaguchi, A. et al. Multistratified Expression of Polysialic Acid and Its Relationship to VAChT-containing Neurons in the Inner Plexiform Layer of Adult Rat Retina. J. Histochem. Cytochem. 47, 919–927 (1999).

61. Fliesler, S. J., Cole, G. J. & Adler, A. J. Neural cell adhesion molecule (NCAM) in adult vertebrate retinas: Tissue localization and evidence against its role in retina-pigment epithelium adhesion. Exp. Eye Res. 50, 475–482 (1990).

62. Becker, C. G., Becker, T., Schmidt, A. & Roth, G. Polysialic acid expression in the salamander retina is inducible by thyroxine. Dev. Brain Res. 79, 140–146 (1994).

63. Kustermann, S., Hildebrandt, H., Bolz, S., Dengler, K. & Kohler, K. Genesis of rods in the zebrafish retina occurs in a microenvironment provided by polysialic acid-expressing Müller glia. J. Comp. Neurol. 518, 636–646 (2010).

64. Fiorito, G. et al. Guidelines for the Care and Welfare of Cephalopods in Research-A consensus based on an initiative by CephRes, FELASA and the Boyd Group. Lab. Anim. 49, 1–90 (2015).

65. Enterina, J. R. et al. Coordinated changes in glycosylation regulate the germinal center through CD22. Cell Rep. 38, 110512 (2022).

66. Morris, R. J. Thy-1, a Pathfinder Protein for the Post-genomic Era. Front. Cell Dev. Biol. 6, 173 (2018).

67. Hagood, J. S. Thy-1 as an Integrator of Diverse Extracellular Signals. Front. Cell Dev. Biol. 7, 26 (2019).

68. Barnstable, C. J. & Dräger, U. C. Thy-1 antigen: A ganglion cell specific marker in rodent retina. Neuroscience 11, 847–855 (1984).

69. Raymond, I. D., Vila, A., Huynh, U.-C. N. & Brecha, N. C. Cyan fluorescent protein expression in ganglion and amacrine cells in a thy1-CFP transgenic mouse retina. Mol. Vis. 14, 1559–1574 (2008).

70. Partida, G. J., Stradleigh, T. W., Ogata, G., Godzdanker, I. & Ishida, A. T. Thy1 Associates with the Cation Channel Subunit HCN4 in Adult Rat Retina. Investig. Opthalmology Vis. Sci. 53, 1696 (2012).

71. Haefliger, J.-A. et al. Targeting Cx40 (Connexin40) Expression or Function Reduces Angiogenesis in the Developing Mouse Retina. Arterioscler. Thromb. Vasc. Biol. 37, 2136–2146 (2017).

72. de Lemos, L. et al. Modelling neurodegeneration and inflammation in early diabetic retinopathy using 3D human retinal organoids. Vitro Models 3, 33–48 (2024).

73. Tajik, A., Phillips, K. L., Nitz, M. & Willis, L. M. A new ELISA assay demonstrates sex differences in the concentration of serum polysialic acid. Anal. Biochem. 600, 113743 (2020).

74. Mowiol mounting medium. Cold Spring Harb. Protoc. 2006, pdb.rec10255 (2006).

75. Schindelin, J. et al. Fiji[]: an open-source platform for biological-image analysis. 9, (2019).

76. Jokilammi, A. et al. Construction of antibody mimics from a noncatalytic enzyme–detection of polysialic acid. J. Immunol. Methods 295, 149–160 (2004).

77. Rutishauser, U., Watanabe, M., Silver, J., Troy, F. A. & Vimr, E. R. Specific alteration of NCAM-mediated cell adhesion by an endoneuraminidase. J. Cell Biol. 101, 1842–1849 (1985).

78. Pelkonen, S., Aalto, J. & Finne, J. Differential activities of bacteriophage depolymerase on bacterial polysaccharide: binding is essential but degradation is inhibitory in phage infection of K1-defective Escherichia coli. J. Bacteriol. 174, 7757–7761 (1992).

79. Hunter, C. et al. Site-specific immobilization of the endosialidase reveals QSOX2 is a novel polysialylated protein. Glycobiology 34, cwae026 (2024).

80. Yu, C.-C. et al. A glyco-gold nanoparticle based assay for α-2,8-polysialyltransferase from Neisseria meningitidis. Chem. Commun. 49, 10166 (2013).

81. Bitter-Suermann, D. & Roth, J. Monoclonal antibodies to polysialic acid reveal epitope sharing between invasive pathogenic bacteria, differentiating cells and tumor cells. Immunol. Res. 6, 225–237 (1987).

82. Frosch, M., Görgen, I., Boulnois, G. J., Timmis, K. N. & Bitter-Suermann, D. NZB mouse system for production of monoclonal antibodies to weak bacterial antigens: isolation of an IgG antibody to the polysaccharide capsules of Escherichia coli K1 and group B meningococci. Proc. Natl. Acad. Sci. 82, 1194–1198 (1985).

83. Kimmel, C. B., Ballard, W. W., Kimmel, S. R., Ullmann, B. & Schilling, T. F. Stages of embryonic development of the zebrafish. Dev. Dyn. 203, 253–310 (1995).

84. Api, M. et al. Effects of Parental Aging During Embryo Development and Adult Life: The Case of Nothobranchius furzeri. Zebrafish 15, 112–123 (2018).

85. Nieuwkoop, P. D. & Faber, J. Normal table of Xenopus laevis (Daudin): a systematical and chronological survey of the development from the fertilized egg till the end of metamorphosis. (1994).

86. Lenkowski, J. R. & Raymond, P. A. Müller glia: Stem cells for generation and regeneration of retinal neurons in teleost fish. Prog. Retin. Eye Res. 40, 94–123 (2014).

87. Singh, R. K., Kolandaivelu, S. & Ramamurthy, V. Early Alteration of Retinal Neurons in Aipl1^−/−^ Animals. Investig. Opthalmology Vis. Sci. 55, 3081 (2014).

88. Daniloff, J., Chuong, C., Levi, G. & Edelman, G. Differential distribution of cell adhesion molecules during histogenesis of the chick nervous system. J. Neurosci. 6, 739–758 (1986).

89. Saito, M., Kitamura, H. & Sugiyama, K. Occurrence of gangliosides in the common squid and pacific octopus among protostomia. Biochim. Biophys. Acta BBA - Biomembr. 1511, 271–280 (2001).

90. Zuber, C., Lackie, P. M., Catterall, W. A. & Roth, J. Polysialic acid is associated with sodium channels and the neural cell adhesion molecule N-CAM in adult rat brain. J. Biol. Chem. 267, 9965–9971 (1992).

91. Galuska, S. P. et al. Synaptic cell adhesion molecule SynCAM 1 is a target for polysialylation in postnatal mouse brain. Proc. Natl. Acad. Sci. U. S. A. 107, 10250–10255 (2010).

92. Curreli, S., Arany, Z., Gerardy-Schahn, R., Mann, D. & Stamatos, N. M. Polysialylated neuropilin-2 is expressed on the surface of human dendritic cells and modulates dendritic cell-T lymphocyte interactions. J. Biol. Chem. 282, 30346–30356 (2007).

93. Yabe, U., Sato, C., Matsuda, T. & Kitajima, K. Polysialic acid in human milk. CD36 is a new member of mammalian polysialic acid-containing glycoprotein. J. Biol. Chem. 278, 13875–13880 (2003).

94. Ribic, A., Liu, X., Crair, M. C. & Biederer, T. Structural organization and function of mouse photoreceptor ribbon synapses involve the immunoglobulin protein synaptic cell adhesion molecule 1. J. Comp. Neurol. 522, 900–920 (2014).

95. Fujita, E., Urase, K., Soyama, A., Kouroku, Y. & Momoi, T. Distribution of RA175/TSLC1/SynCAM, a member of the immunoglobulin superfamily, in the developing nervous system. Dev. Brain Res. 154, 199–209 (2005).

96. Pietri, T., Easley-Neal, C., Wilson, C. & Washbourne, P. Six cadm/synCAM genes are expressed in the nervous system of developing zebrafish. Dev. Dyn. 237, 233–246 (2008).

97. Wang, J., Ou, S.-W. & Wang, Y.-J. Distribution and function of voltage-gated sodium channels in the nervous system. Channels Austin Tex 11, 534–554 (2017).

98. Kiss, J. Z., Troncoso, E., Djebbara, Z., Vutskits, L. & Muller, D. The role of neural cell adhesion molecules in plasticity and repair. Brain Res. Rev. 36, 175–184 (2001).

99. Ono, K., Tomasiewicz, H., Magnuson, T. & Rutishauser, U. N-CAM mutation inhibits tangential neuronal migration and is phenocopied by enzymatic removal of polysialic acid. Neuron 13, 595–609 (1994).

100. Silver, J. & Rutishauser, U. Guidance of optic axons in vivo by a preformed adhesive pathway on neuroepithelial endfeet. Dev. Biol. 106, 485–499 (1984).

101. Tang, J., Landmesser, L. & Rutishauser, U. Polysialic acid influences specific pathfinding by avian motoneurons. Neuron 8, 1031–1044 (1992).

102. Yin, X., Watanabe, M. & Rutishauser, U. Effect of polysialic acid on the behavior of retinal ganglion cell axons during growth into the optic tract and tectum. Development 121, 3439–3446 (1995).

103. Muller, D. et al. PSA–NCAM Is Required for Activity-Induced Synaptic Plasticity. Neuron 17, 413–422 (1996).

104. Muller, D., Stoppini, L., Wang, C. & Kiss, J. Z. A role for polysialylated neural cell adhesion molcule in lesion-induced sprouting in hippocampal organotypic cultures. Neuroscience 61, 441–445 (1994).

105. Aubert, I., Ridet, J. L., Schachner, M., Rougon, G. & Gage, F. H. Expression of L1 and PSA during sprouting and regeneration in the adult hippocampal formation. J. Comp. Neurol. 399, 1–19 (1998).

106. Noel, N. C. L., MacDonald, I. M. & Allison, W. T. Zebrafish Models of Photoreceptor Dysfunction and Degeneration. Biomolecules 11, 78 (2021).

107. Bergmans, S. et al. Age-related dysregulation of the retinal transcriptome in African turquoise killifish. Aging Cell 23, e14192 (2024).

108. Vanhunsel, S. et al. The killifish visual system as an in vivo model to study brain aging and rejuvenation. Npj Aging Mech. Dis. 7, 22 (2021).

109. Noel, N. C. et al. Characterising Photoreceptor Subtypes in the Turquoise Killifish (Nothobranchius furzeri). Invest. Ophthalmol. Vis. Sci. 64, 3735–3735 (2023).

110. Distler, C. & Dreher, Z. Glia Cells of the Monkey Retina—II. Müller Cells. Vision Res. 36, 2381–2394 (1996).

111. Dreher, Z., Robinson, S. R. & Distler, C. Müller cells in vascular and avascular retinae: A survey of seven mammals. J. Comp. Neurol. 323, 59–80 (1992).

112. Dreher, Z., Distler, C. & Dreher, B. Vitread proliferation of filamentous processes in avian Müller cells and its putative functional correlates. J. Comp. Neurol. 350, 96–108 (1994).

113. Masland, R. H. The fundamental plan of the retina. Nat. Neurosci. 4, 877–886 (2001).

114. Haverkamp, S. & Wässle, H. Immunocytochemical analysis of the mouse retina. J. Comp. Neurol. 424, 1–23 (2000).

115. Brunk, U. T. & Terman, A. Lipofuscin: mechanisms of age-related accumulation and influence on cell function. Free Radic. Biol. Med. 33, 611–619 (2002).

116. Călin, E. F. et al. Lipofuscin: a key compound in ophthalmic practice. Romanian J. Ophthalmol. 65, 109–113 (2021).

117. Forestier, N. L., Lescs, M. & Gherardi, R. K. Anti-NKH-1 antibody specifically stains unmyelinated fibres and non-myelinating Schwann cell columns in humans. Neuropathol. Appl. Neurobiol. 19, 500–506 (1993).

118. Bosch-Queralt, M., Fledrich, R. & Stassart, R. M. Schwann cell functions in peripheral nerve development and repair. Neurobiol. Dis. 176, 105952 (2023).

119. Lashay, A. et al. Role of Schwann Cells in Preservation of Retinal Tissue Through Reduction of Oxidative Stress. Med. Hypothesis Discov. Innov. Ophthalmol. J. 8, 323–332 (2019).

120. Dezawa, M. & Adachi-Usami, E. Role of Schwann cells in retinal ganglion cell axon regeneration. Prog. Retin. Eye Res. 19, 171–204 (2000).

121. Garcia-Garcia, D., Locker, M. & Perron, M. Update on Müller glia regenerative potential for retinal repair. Curr. Opin. Genet. Dev. 64, 52–59 (2020).

122. Poonam, S. & Rajesh, R. Retina regeneration: lessons from vertebrates. Oxf. Open Neurosci. 1–21 (2022).

123. Bu, Q. et al. Neurodevelopmental defects in human cortical organoids with N-acetylneuraminic acid synthase mutation. Sci. Adv. 9, (2023).

124. Krishnan, A. et al. PolySialic acid-nanoparticles inhibit macrophage mediated inflammation through Siglec agonism: a potential treatment for age related macular degeneration. Front. Immunol. 14, 1237016 (2023).

125. Karlstetter, M. et al. Polysialic acid blocks mononuclear phagocyte reactivity, inhibits complement activation, and protects from vascular damage in the retina. EMBO Mol. Med. 9, 154–166 (2017).

126. Lobanovskaya, N. & Zharkovsky, A. A role of PSA-NCAM in the survival of retinal ganglion cells (RGCs) after kainic acid damage. NeuroToxicology 72, 101–106 (2019).

127. Martinez-De Luna, R. I., et al. Müller glia reactivity follows retinal injury despite the absence of the glial fibrillary acidic protein gene in Xenopus. Dev. Biol. 426, 219–235 (2017).

128. Gorovits, R., Avidan, N., Avisar, N., Shaked, I. & Vardimon, L. Glutamine synthetase protects against neuronal degeneration in injured retinal tissue. Proc. Natl. Acad. Sci. 94, 7024–7029 (1997).

129. Lewis, G. P. & Fisher, S. K. Up-Regulation of Glial Fibrillary Acidic Protein in Response to Retinal Injury: Its Potential Role in Glial Remodeling and a Comparison to Vimentin Expression. in International Review of Cytology vol. 230 263–290 (Elsevier, 2003).

