## Supplemental Data for "Polysialic acid is a versatile marker for retinal Müller glia in common vertebrate model organisms and systems"

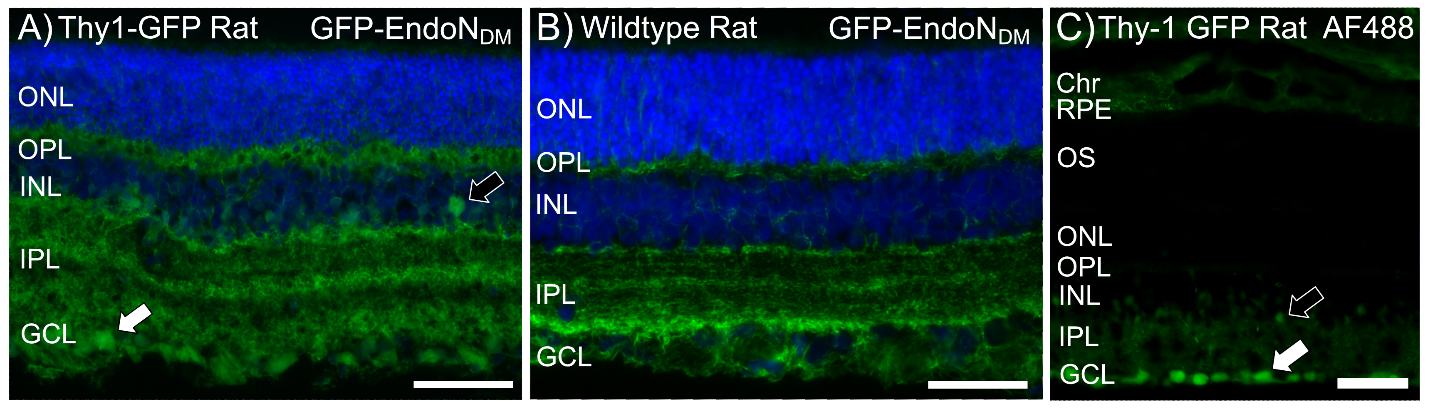

**Supplementary Figure S1.** Comparison of Thy-1 GFP+ polySia immunoreactivity, Thy-1 GFP + AF488 non-specific cross reactivity in Thy-1 GFP transgenic rats, and polySia immunoreactivity in wildtype rats. There was additional GFP+ signal in a subset of amacrine cells (black arrows) and ganglion cells (white arrows) in the transgenic rats (A) compared to wildtype rats (B), but there was no significant difference in polySia expression patterns. AF488 non-specific cross reactivity was apparent in the RPE and choroid of Thy-1 GFP transgenic rats (C), and the amacrine cell and ganglion cell GFP positive signal from the Thy-1 GFP was also apparent (C). *Labels*: Green, GFP-EndoN_DM_(A,B), AF448/GFP (C); Blue, Hoechst. *Abbreviations*: Chr, choroid; RPE, retinal pigment epithelium; OS, outer segments; ONL, outer nuclear layer; OPL, outer plexiform layer; INL, inner nuclear layer; IPL, inner plexiform layer; GCL, ganglion cell layer. Scale bar = 50 µm. Thy1-GFP n = 2, wildtype n = 1.

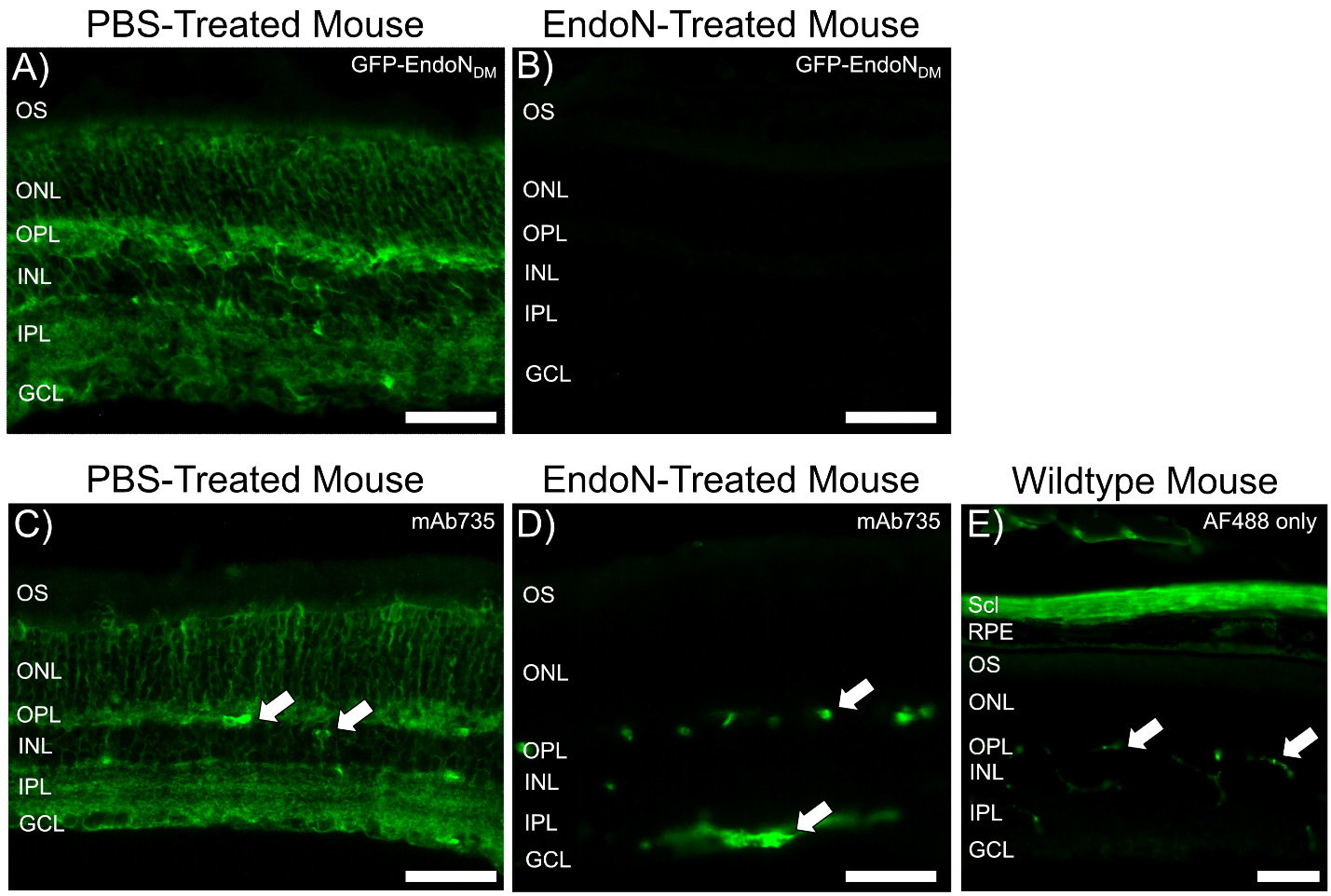

**Supplementary Figure S2**. Mouse retinal tissues pre-incubated with PBS (A,C) and endoneuraminidase (B,D). GFP-EndoN and mAb735 reactivity is lost when sections are pre-treated with polySia hydrolase, which cleaves polySia chains. Cross-reactivity with blood plasma IgGs in intra-retinal and vitreal blood vessels (white arrows) is seen due to the use of the anti-mouse secondary antibody for mAb735 detection (E). *Labels*: Green, GFP-EndoN_DM_ (A), mAb735 (C), or AF488 only (E). *Abbreviations*: OS, outer segments; ONL, outer nuclear layer; OPL, outer plexiform layer; INL, inner nuclear layer; IPL, inner plexiform layer; GCL, ganglion cell layer. Scale bar = 50 µm. n = 3.

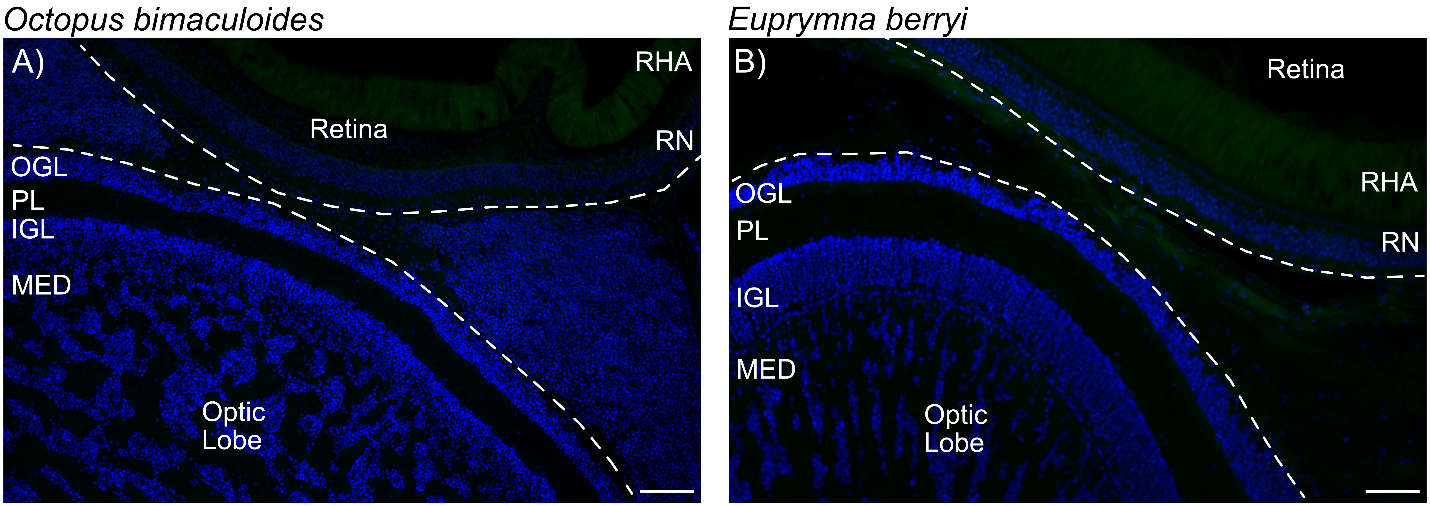

**Supplementary Figure S3.** GFP-EndoN_DM_ does not label cephalopod retina or optic lobe. There was no significant labeling in the octopus (A) or squid (B) retina or optic lobe using GFP-EndoN_DM_. *Labels*: Green, mAb735; Blue, Hoechst. *Abbreviations*: RHA, rhabdomeres; RN, retinal nuclei; OGL, outer granular layer; PL, plexiform layer; IGL, inner granular layer; MED, medulla. Scale bar = 50 µm. n = 1.

**Supplemental Table S1.** Antigens, species/type, source, and working dilutions of antibodies used in this study.

| **Antigen** | **Species/Type** | **Source** | **Concentration** |
| --- | --- | --- | --- |
| **Primary Antibodies & Lectins** | | | |
| GFP-EndoN_DM_ | lectin | L.M. Willis^[1]^ | 10 µg/mL |
| mAb735  (Clone 735) | mouse monoclonal | Absolute Antibody  AB00240 | 1 µg/mL |
| NCAM  (Clone 4d) | mouse monoclonal | DHSB^[2]^ | 0.7 µg/mL |
| GS  (Clone GS-6) | mouse monoclonal | Millipore Sigma  MAB302 | 2 µg/mL |
| Vimentin  (Clone 14h7) | mouse monoclonal | DHSB^[3]^ | 7.6 µg/mL |
| GFAP  (Clone GA5) | mouse monoclonal | Millipore Sigma  G3893 | 0.1 mg/mL |
| **Secondary Antibodies and Nuclear Stains** | | | |
| Hoechst 33342 | nuclear stain | Sigma Aldrich | 0.1 mg/mL |
| Cy3 | anti-mouse | Jackson ImmunoResearch | 2 µg/mL |

1. Tajik, A., Phillips, K. L., Nitz, M. & Willis, L. M. A new ELISA assay demonstrates sex differences in the concentration of serum polysialic acid. Anal. Biochem. 600, 113743 (2020).
2. Developmental Studies Hybridoma Bank; deposited by Rutishauser, U.
3. Developmental Studies Hybridoma Bank; deposited by Klymkowsky, M.
